# Uncovering the Structural and Functional Implications of Uncharacterized NSPs and Variations in the Molecular Toolkit Across Mammalian and Non-Mammalian Arteriviruses

**DOI:** 10.1101/2024.12.01.626221

**Authors:** Rakesh Siuli, Kshitij Behera, Arunkumar Krishnan

**Author notes:** **Author for correspondence** Krishnan A. Email IDs: Rakesh S, Kshitij B.

## Abstract

Despite considerable scrutiny of mammalian arterivirus genomes, their genomic architecture remains incomplete, with several unannotated non-structural proteins (NSPs) and the puzzling absence of methyltransferase (MTase) domains. Additionally, the host range of arteriviruses has expanded to include seven newly sequenced genomes from non-mammalian hosts, which remain largely unannotated and await detailed comparisons alongside mammalian isolates. Utilizing comparative genomics approaches and comprehensive sequence-structure analysis, we provide enhanced genomic architecture and annotations for arterivirus genomes. We identify the previously unannotated C-terminal domain of NSP3 as a winged helix-turn-helix domain and classified NSP7 as a new small β-barrel domain, both likely involved in interactions with viral RNA. NSP12 is identified as a derived variant of the N7-MTase-like Rossmann fold domain, showing structural alignment with N7-MTases in Nidovirales, yet it likely lacks enzymatic functionality due to the erosion of catalytic residues, indicating a unique role-specific to mammalian arteriviruses. In contrast, non-mammalian arteriviruses sporadically retain a 2ʹ O-MTase and an exonuclease domain, which are typically absent in mammalian arteriviruses, highlighting contrasting evolutionary trends and variations in their molecular armory. Similar lineage-specific paterns are observed in the diversification of papain-like proteases and structural proteins. Overall, the study extends our knowledge of arterivirus genomic diversity and evolution.

## Introduction

Arteriviruses are positive-sense, single-stranded RNA viruses with relatively small genomes and distinct virion structures, belonging to the order Nidovirales (**1–3**). They predominantly infect boreoeutherian mammals such as horses, pigs, rodents, and non-human primates, causing critical illnesses like hemorrhagic fever and severe respiratory diseases, particularly in porcine and equine hosts (**4–6**). Their high virulence and contagious nature pose significant challenges to the porcine and equine industries, leading to extensive research into their various molecular aspects to identify druggable targets and develop effective control measures. Within the Nidovirales order, arteriviruses have followed a distinct evolutionary path, establishing themselves as one of the four primary families alongside Mesoniviridae, Roniviridae, and Coronaviridae (**3,7,8**). This unique trajectory positions arteriviruses as an essential model for investigating the broader evolutionary paterns, especially in comparison to other Nidovirale families.

Arterivirus genomes typically range from 13 to 15 kb in size, consisting of 10 to 15 ORFs, with the two longest ORFs, ORF1a and ORF1b, positioned sequentially from the 5’ end, and encoding a series of non-structural proteins (NSPs) that are crucial for viral genome replication. These ORFs are translated into two large replicase polyproteins, polyprotein1a (pp1a) and polyprotein1b (pp1b). Current knowledge suggests that the genomic structure of ORF1a is characterized by one or more papain-like proteases (PLPs), followed by a series of transmembrane (TM) segments and a 3C-like protease (3CL-Pro). Upon translation, PLPs process the ORF1a encoded pp1a polyprotein into seven NSPs and thereby facilitate the maturation and assembly of the replication-transcription complex (RTC) (**9–11**). While PLPs process ORF1a into seven NSPs, the 3CL-pro (NSP4) encoded in ORF1a primarily cleaves and processes downstream NSPs within ORF1b into four NSPs (NSP9 to NSP12). These NSPs are key for RNA replication and processing (**12,13**) and include the following: the RNA-dependent RNA polymerase (RdRp) in NSP9 (**14**), a zinc-binding and helicase domain in NSP10 (**15**), an endoribonuclease domain (NendoU) in NSP11 (**16**), and finally, NSP12, which encodes a globular domain (**17**), whose structural and functional roles remain largely unexplored. Further downstream of ORF1b, a variable number of shorter ORFs encode various structural proteins, including viral membrane (M) proteins, nucleocapsid (N) proteins, and multiple envelope glycoproteins (GP) that aid in virion assembly and release from the host cell (**18,19**). These minor ORFs exhibit considerable variation in both number and sequence-structure synapomorphies, with some likely arising from ancient duplication events and subsequently following distinct evolutionary trajectories shaped by viral-host coevolution (**20**), contributing to the overall diversity in arterivirus genome size.

Despite notable research progress on arteriviruses from mammalian hosts (hereafter mammalian arteriviruses), there are still significant gaps in our understanding of their complete genomic architecture and protein products. While many NSPs and their constituent domains have been studied, others, such as NSP3C (the C-terminal end of NSP3), NSP7 (within ORF1a), and NSP12 (C-terminal end of ORF1b), remain poorly understood and insufficiently characterized. This lack of detailed structural and functional information on these elusive ORFs impedes a complete understanding of arterivirus genomic architecture and their functional implications. Moreover, in 2018, Shi Mang, et al. published 214 vertebrate-associated RNA viral genomes from a wide range of vertebrate hosts, including several from non-mammalian sources, for the first time. These included six different arterivirus genomes: one from ray-finned fish, one from cartilaginous fish, and four from reptilian (three snakes and one turtle) hosts (**21**), all of which remain unannotated and largely unstudied. These genomes, along with an independently sequenced arterivirus genome from a reptilian host (Chinese softshell turtle; *Trionyx sinensis* hemorrhagic syndrome arterivirus -TSHSA) (**22**), formed the core set of seven arterivirus genomes from non-mammalian hosts (referred to as non-mammalian arteriviruses hereafter) that we analyzed and compared with mammalian arteriviruses for the first time in our study. For mammalian arteriviruses, we compiled a set of 22 full-length arterivirus genomes from the NCBI RefSeq database, covering all major mammalian host groups (primates, porcine, rodents, equine, and marsupials). This inclusive set of arteriviruses from both mammalian and non-mammalian hosts enabled us to analyze complete protein domain compositions, annotate previously uncharacterized segments, and perform an in-depth comparative genomics analysis to illuminate broader aspects of arterivirus evolution and diversification.

In this study, we present the first comprehensive annotation of non-mammalian arteriviruses, including the exclusive identification of SAM-dependent 2’-O-MTase and ExoN (Exonuclease) domains in these viruses. The analysis uncovered significant differences between arteriviruses from mammalian and non-mammalian hosts, with the latter exhibiting greater variation in genome size, ORF count, and domain composition. Additionally, the study presents the initial characterization of NSP3C and NSP7 within ORF1a in mammalian arteriviruses, linking them to well-known nucleic acid binding domains, and demonstrated that NSP12 represents a divergent form of Rossmann fold containing MTase-like domain. Additionally, we identified host-specific divergence in the glycoproteins of mammalian arteriviruses and uncovered several glycoproteins in newly sequenced non-mammalian arteriviruses, providing comparative insights into these structural proteins. Through detailed comparative genomics and in-depth sequence-structure analysis, we systematically classified several long-overlooked and previously undefined functional domains, deepening our understanding of arterivirus evolution and biology.

## Materials and Methods

### Genomic datasets and sequence analysis

To facilitate a comprehensive comparison of arterivirus genomes from both mammalian and non-mammalian hosts, we curated a dataset of 22 complete mammalian arterivirus genomes from the NCBI RefSeq database, excluding those not listed in RefSeq. These genomes encompass a range of boreoeutherian mammals, including both primates and non-primates. Alongside the mammalian genomes, we incorporated all currently available non-mammalian arterivirus genome sequences. This includes genomes from snakes, turtles, ray-finned fish (Japanese halfbeak), and cartilaginous fish (Ghost shark). All encoded protein FASTA sequences from these genomes were obtained and clustered using the BLASTCLUST program (ftp://ftp.ncbi.nih.gov/blast/documents/blastclust.html) (<u>RRID:SCR_016641</u>; version 2.2.26). The parameters, including the length of pairwise alignments (L) and the bit-score (S), were adjusted to achieve the desired level of clustering by empirically modifying the alignment length and bit-score density threshold. Divergent sequences or smaller clusters were merged with larger ones when supported by supplementary evidence, including shared sequence motifs, structural synapomorphies, reciprocal BLAST search results, and/or associations in genome context. To accurately define the encoded protein domains and their boundaries for each ORF, sequences from all analyzed genomes were subjected to sensitive profile-profile searches using HHPRED against HMM (Hidden Markov Model) profiles derived from the PDB (**23**) and Pfam models (**24**). For each query or seed sequence analyzed through HHPRED, HHblits was utilized with default parameters to retrieve homologs from the UniRef30 database, subsequently creating Multiple Sequence Alignments (MSAs) and corresponding HMM models for profile-profile comparison (**25–27**). Additionally, RPSBLAST (**28**) with default parameters was used to perform searches against our custom in-house database of diverse domains. MSAs for all domains analyzed in this study were generated using MAFFT (**29**) or Kalign (V3) (**30**), with manual adjustments based on profile-profile searches against PDB, structural alignments, and predicted 3D structural models (see below). Transmembrane regions were predicted using Deep TMHMM with default settings (**31**).

### Structure analysis

3D structure predictions for all individual domains analyzed in this study were generated using AF3 (**32**). Each predicted structure was then subjected to structural similarity searches with the DALI-lite program (RRID:SCR_003047) (**33**) against the PDB database clustered at 75% identity. The reference MSAs constructed for each analyzed domain were used to predict secondary structure topologies with the JPred (V4) program (RRID:SCR_016504) (**34**), and the predicted secondary structure boundaries from JPred4 were cross-validated with the AF3 models. Structural homology among domains was evaluated based on DALI Z-scores and 3D structural superimposition, with manual validation for accuracy. Structures were rendered, compared, and superimposed using PyMOL (https://www.pymol.org/) (<u>RRID: SCR_000305</u>).

### Comparative genomics and phylogenetic analysis

Structural glycoproteins of arteriviruses from both mammalian and non-mammalian hosts were clustered by calculating all-against-all pairwise sequence similarities using the CLANS software (**35**). Host-specific divergence and phylogenetic relationships of each structural glycoprotein were inferred with an approximate maximum likelihood (ML) approach in the FastTree program (RRID: SCR_015501) (**36**), with local support values estimated accordingly. To enhance the topology’s accuracy, the number of minimum-evolution subtree-prune-regraft (SPR) rounds in FastTree was set to 4 (-spr 4), and the options ‘-mlacc’ and ‘-slownni’ were applied for more exhaustive ML nearest neighbor interchanges (NNIs). Phylogenetic tree topologies were also generated using ML methods based on the edge-linked partition model in the IQ-TREE software (**37,38**), with branch support obtained via the ultrafast bootstrap method (1000 replicates) in IQ-TREE (**39**). The FigTree program (RRID:SCR_008515) (http://tree.bio.ed.ac.uk/software/figtree/) was used to render phylogenetic trees.

## Results and Discussion

### Evolutionary diversification of N-terminal PLPs and distinguishing features of newly characterized domains in ORF1a

In mammalian arteriviruses, the N-terminal region of ORF1a is characterized by several PLPs, with NSP1 encoding two to three of these proteases—PLP1α, PLP1β, and PLP1γ (**Figure 1A**). PLP1α and PLP1β are conserved across all mammalian arteriviruses, while PLP1γ is specific to primate arteriviruses. In addition, NSP2 encodes a conserved PLP, known as PLP2, which is recognized for its deubiquitinase activity that interferes with host immune proteins (**11,40,41**). These PLP paralogs were earlier implied to have likely emerged via early duplication events followed by diversifying positive selection, leading to their structural and functional diversifications (**41**). In non-mammalian arteriviruses, we surprisingly found a marked reduction in the number of PLPs within ORF1a compared to their mammalian counterparts (**Figure 1B**). Both PLP1β and PLP1γ are absent, and PLP1α is retained solely in the genome of Hainan *Oligodon formosanus* arterivirus (HOFA - Reptilia). Notably, PLP2 was found in all analyzed non-mammalian arterivirus genomes except for *Nanhai ghost shark* arterivirus (NGSA – Chondrichthyes) and Hainan *Hebius popei* arterivirus (HHPA - Reptilia) (**Figure 1B**). Sequence-structure analysis reveals that these non-mammalian PLPs, particularly PLP2, subtly differ from their mammalian analogs, exhibiting slightly distinct features. Nevertheless, all five PLP2s share a conserved catalytic motif, DGxCGhH (**41**) in the first α-helix, as well as a conserved catalytic histidine residue in the fourth β-strand, which are also present in all PLP2s across the arteriviruses analyzed (**Supplementary Datasets S1**).

**Figure 1.**
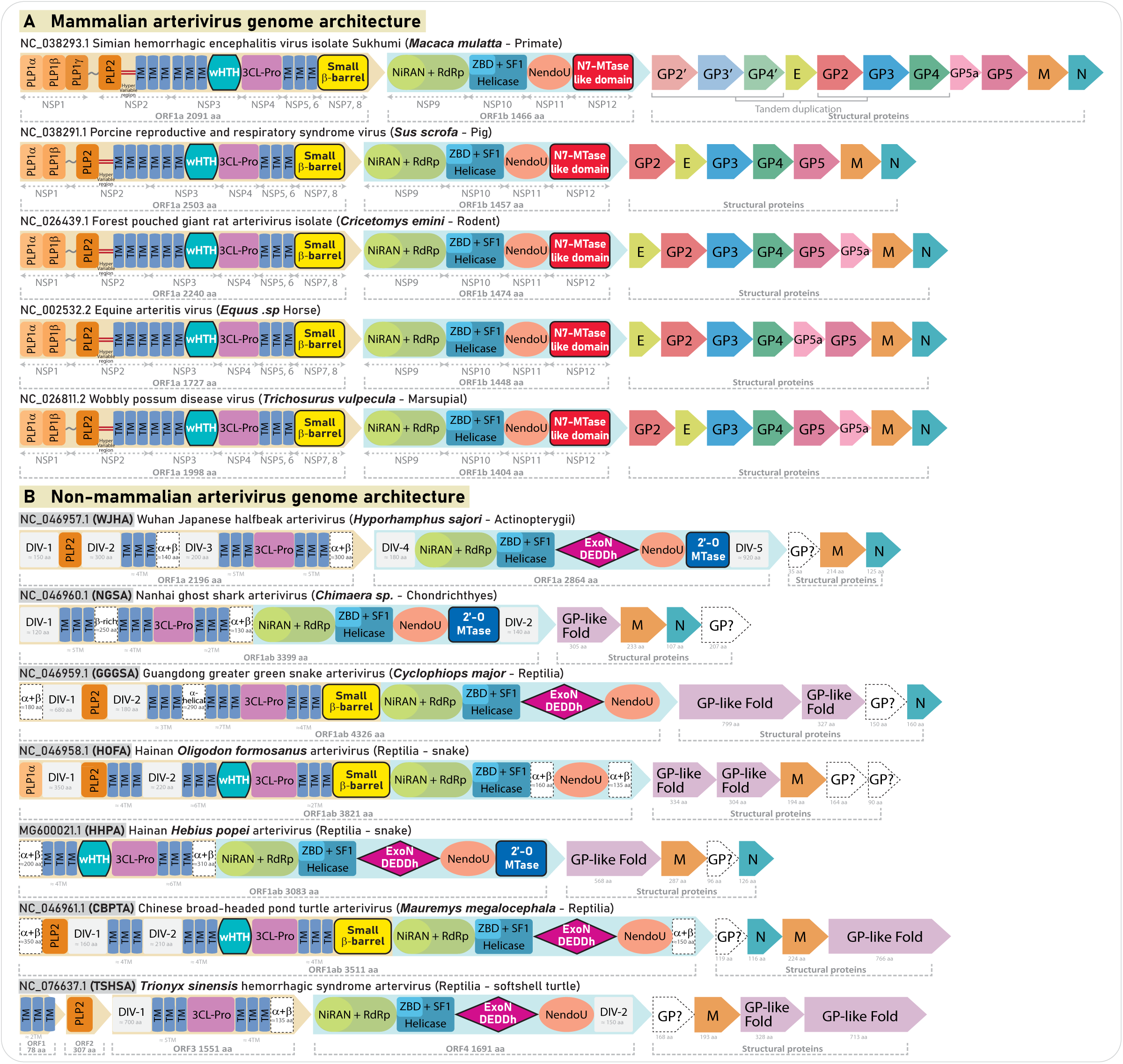
Genomic architecture and proteome composition of arterivirus genomes. **(A)** Genomic structure and protein domain organization of representative mammalian arteriviruses. ORFs are depicted as colored arrows, with their encoded domains labeled and highlighted. The color scheme is consistently applied across the figure for homologous regions and domains. Newly identified functional domains—the C-terminal wHTH domain of NSP3, the small β-barrel domain of NSP7, and the N7-MTase-like domain of NSP12—are outlined with black borders **(B)** Genomic structure and protein domain organization of arterivirus genomes from non-mammalian hosts. In cases where ORF1a and ORF1b regions are merged into a single, larger ORF (as observed in NGSA, GGGSA, HOFA, HHPA, and CBPTA genomes) or split into multiple shorter ORFs (as in TSHSA genome), the corresponding regions of ORF1a and ORF1b are highlighted using the same color scheme as their mammalian counterparts (light brown background for ORF1a and light blue for ORF1b) to aid easier comparison. Highly divergent genomic regions predicted to be largely disordered with few minor secondary structural elements are marked as DIVs with grey background. Likely functional domains with predicted secondary structural elements and ordered structures, as inferred from AF3, but not yet annotated in ORF1a and ORF1b due to a lack of homology with known domains, are displayed with a white background and dotted borders. Similarly, GP-like domains that lack clear structural homology for precise annotation are labeled as GP? and shown with a white background and dotted borders. Newly identified and classified functional domains, including the wHTH, small β-barrel, 2ʹ-O-MTase, and ExoN domains, are outlined in black. Each genome shown in both panels is labeled with its NCBI RefSeq accession number, species name, common name. Abbreviations are provided for non-mammalian genomes for easy usage in the manuscript.

Though no additional PLPs were found, most non-mammalian arteriviruses have at least one α+β globular domain, typically located at the N-terminus or in genomic locations analogous to NSP1/NSP2-like domains found in mammalian arteriviruses (**Figure 1B, Supplementary Figure S1**). Predictions from AlphaFold 3 (AF3) suggest αβα-sandwich-like architecture for most of these domains, however, no clear homology to known proteins were observed in DALI structure similarity searches. Besides these segments, several of these genomes encode highly divergent intervening regions within the ORF1a, typically 120–300 amino acids, with some extending beyond 400–600 residues, as in TSHSA (**Figure 1B**). Sequence and structural analysis revealed no associations with PLPs or other functional domains, with most regions being unalignable. AF3 predictions indicate these regions are predominantly disordered, with only a few minor segments demonstrating ordered secondary structure elements such as beta-hairpins and helical bundles embedded within the largely disordered structure. Additionally, no transmembrane segments were identified by Deep-TMHMM in these regions. Collectively we annotate these regions as DIVs (Disordered Intervening Regions) (**Figure 1B, Supplementary Figure S2**). While disordered regions may gain structured configurations during protein-protein interactions, the potential roles of these DIVs remain uncertain. Furthermore, it is unclear whether any of the identified α+β domains could serve enzymatic functions akin to PLPs. Improved genome assemblies and sequencing additional non-mammalian arteriviruses, and experimental characterization of these predicted domains would help clarify these uncertainties.

In addition, ORF1a encodes several transmembrane segments within NSP2 and NSP3, and the C-terminal end of NSP3 (NSP3C) encodes a globular domain that we have now categorized as a winged helix-turn-helix (wHTH) domain (detailed in the next section) (**Figure 1A**). Following NSP3, NSP4 encodes the conserved 3CL-pro in mammalian arteriviruses and these are also consistently retained in similar genomic locations within ORF1a in non-mammalian arteriviruses, preserving their key sequence and structural features (**Figure 1A and 1B**). Non-mammalian arteriviruses also include extra transmembrane segments adjacent to 3CL-pro, mirroring those found in mammals. At the end of the C-terminal region, ORF1a in mammalian arteriviruses encode NSP7, which we have demonstrated to encode a small β-barrel domain (see next sections). Beyond ORF1a, Figure 1 illustrates the complete genomic architecture of arteriviruses, emphasizing all newly identified domains and offering a comprehensive comparative view of the genomic structure in both mammalian and non-mammalian arteriviruses (**Supplementary Table S1**).

### NSP3C is a bona fide wHTH module that likely plays a role in binding single-stranded RNA

Our analysis uncovers that the previously unannotated C-terminal domain of NSP3 in mammalian arteriviruses is a conserved wHTH domain, which is usually defined by two strands and a characteristic tri-helical core with strands forming an anti-parallel β-sheet, functioning as the nucleic-acid binding interface (**42,43**) **(Figure 2A)**. We observe that the wHTH module is conserved across all mammalian arteriviruses and at least three non-mammalian arterivirus genomes (**Figure 1**). Interestingly, when comparing the genomic architecture with other nidovirales, we find that arterivirus NSP3 is equivalent to coronavirus NSP4, both encoding multiple transmembrane (TM) segments (typically 4 to 6) followed by a C-terminal wHTH domain (**44–46**). DALI searches, using crystal structures of coronavirus NSP4C and AF3-predicted arterivirus NSP3C, have revealed multiple structurally overlapping wHTH modules, confirming their homology to canonical wHTH domains. The core structures of NSP3C wHTH in arteriviruses and NSP4C wHTH in coronaviruses share a conserved βS1-αH1-αH2-αH3-βS2 arrangement, with minimal structural deviation observed upon superimposition **(Figure 2A)**. It is known that the TM segments of NSP3, alongside other TM regions of ORF1a, contribute to the formation of double-membrane vesicles (DMVs) that serve as scaffolds for the Replication-Transcription Complex (RTC) (**44,45,47**). In coronaviruses, the TM regions of NSP4 are embedded in the endoplasmic reticulum (ER) derived membrane of DMVs, positioning the C-terminal domain within the DMV (**46,48**). We propose that arterivirus NSP3, with its TM segments and C-terminal wHTH domain, likely serves a similar function, with the wHTH domain interacting specifically with the viral RNA.

**Figure 2.**
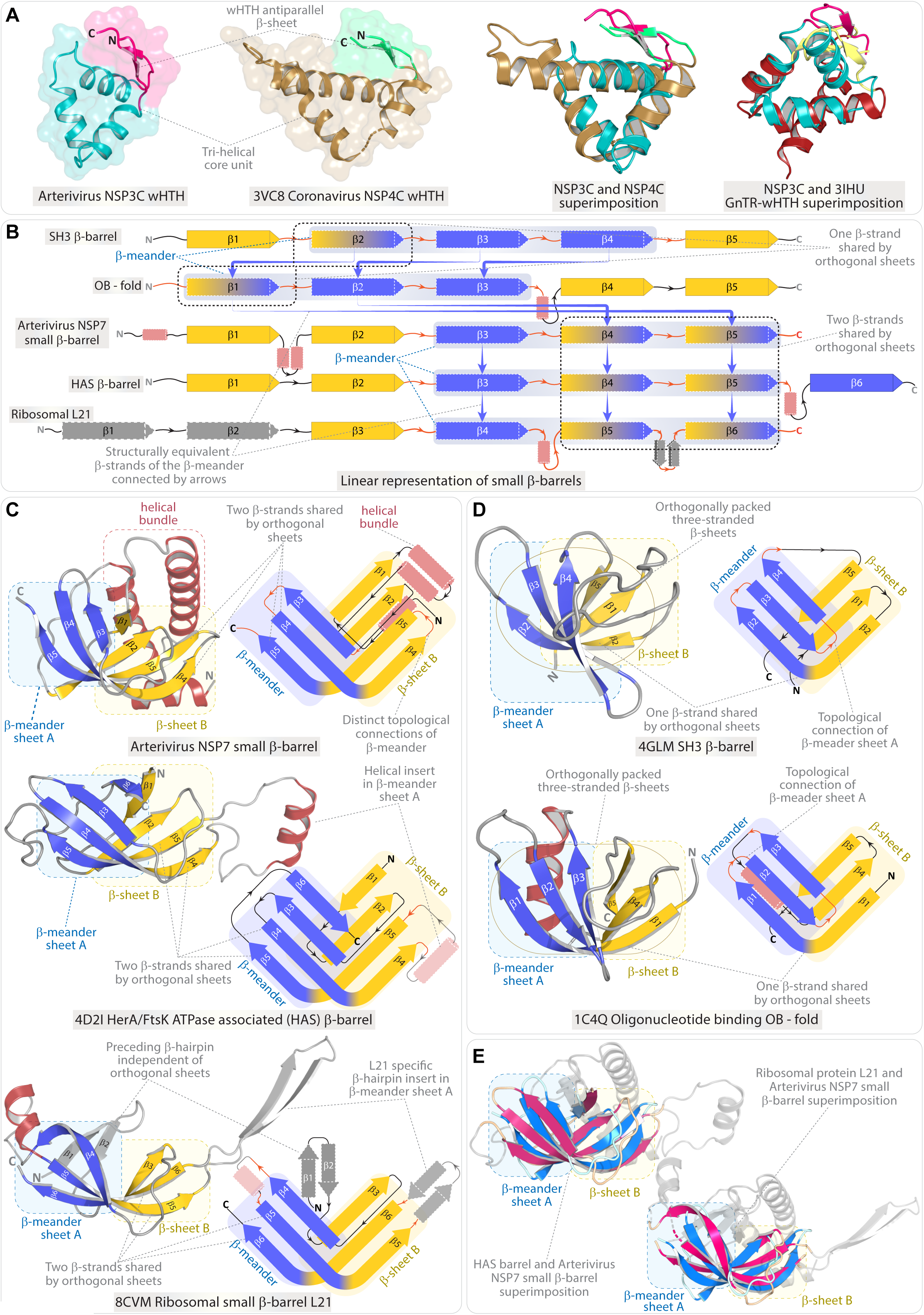
Comparative analysis and structural characteristics of the arterivirus NSP3C wHTH and NSP7 small β-barrel domains. **(A)** The first two 3D structures in panel A illustrate the AF3-predicted C-terminal wHTH domain of NSP3 (NSP3C) from arteriviruses, alongside the experimentally resolved and corresponding wHTH domain in the C-terminal region of NSP4 (NSP4C) from coronaviruses (PDB ID: 3VC8). The structural superpositions on the right display the alignment of NSP3C with NSP4C, as well as NSP3C with the canonical wHTH domain in GnTRs (PDB ID: 3IHU). Panel **(B)** displays a linear topological representation and comparison of the arterivirus NSP7 small β-barrel domain with closely related small β-barrel domains that, while representing distinct folds, share a broader structural framework. These domains are collectively classified within the SBB assemblage, with NSP7 being identified as a new member in this study. **(C)** 3D structure and topological comparisons of the arterivirus NSP7 small β-barrel (AF3-predicted), HerA/FtsK ATPase-associated (HAS) β-barrel (PDB ID: 4D2I), and ribosomal small β-barrel L21 (PDB ID: 8CVM). As shown in panel B, these three domains are closely related and share two β-strands that are common between the nearly orthogonally oriented β-sheet A and β-sheet B, unlike the single shared strand seen in SH3 and OB folds. β-strands forming β-sheet A (β-meander) and β-sheet B are highlighted in light blue and yellow, with the shared strands (common to both sheets) colored accordingly for each half. The schematic topology, displayed alongside the 3D structure, is presented in a simplified format to precisely illustrate the nearly orthogonal β-sheets and the connectivity between the β-strands. **(D)** Representative 3D structures and schematic topology of the SH3-like β-barrel (PDB ID: 4GLM) and oligonucleotide-binding (OB) fold domains (PDB ID: 1C4Q). **(E)** Structural superposition of the arterivirus NSP7 small β-barrel with (i) the HAS barrel and (ii) the ribosomal small β-barrel L21.

### NSP7: a new member of the SBB assemblage with potential roles in viral RNA interaction and nucleoprotein complex stabilization

NSP7, positioned at the 3’-end of ORF1a, encodes a globular domain that remains poorly understood. Like the NSP3C module, this domain is conserved across all mammalian arteriviruses, and we could also identify homologs in at least three non-mammalian arteriviruses at comparable genomic locations (**Figure 1**). Although the 3D structure of NSP7 has been determined using NMR, the protein domain remains unannotated, and its functional role in arteriviruses is still unclear, with previous studies suggesting it may represent a novel fold of unknown function (**49**). Using AF3 predictions, here we demonstrate that all NSP7 domains in mammalian arteriviruses, along with its NSP7-equivalent domain from ORF1a in three non-mammalian arteriviruses (**Figure 1B**), consistently retains a small β-barrel like structural scaffold composed of five core β-strands that assemble into two “nearly” orthogonal and flexible β-sheets, forming a small β-barrel domain. These structural features affiliate with those of previously classified small β-barrel domains (see next paragraph) (**50–52**) (**Figure 2B-D**). Topologically, NSP7 can be depicted as αH1-βS1-(αH2-αH3)-βS2-βS3-βS4-βS5. The preceding helix αH1 and the bihelical hairpin insert region after core βS1 form an N-terminally located helical bundle, which is stacked posterior to the core five-stranded β-barrel (**Figure 2C**).

A recent structure-based survey has classified a broad range of small β-barrel (SBB) domains, incorporating diverse and closely related protein folds that, despite their uniqueness, share a broader structural framework. This allows these domains to be grouped at the ‘superfold’ or ‘urfold’ level (broader structural relationships between multiple closely related protein folds), offering a macroscopic perspective for comparing these SBBs with structural and geometric similarities in their overall β-barrel architecture (**52**). Functionally, these domains also act as diverse nucleic acid-binding moieties and participate in interactions with other proteins (**52,53**). Using sensitive structural similarity searches via DALI, several members of the SBB assemblage were recovered for NSP7, such as the oligonucleotide-binding (OB) fold, small β-barrels in ribosomal proteins, and the HAS barrel domains associated with HerA/FtsK ATPases. Like the NSP7, the members of the SBB group recovered in the DALI searches are usually characterized by a five, or occasionally a six-stranded β-barrel, which can be structurally divided into two orthogonally packed β-sheets—β-sheet A and β-sheet B—that share at least one β-strand (**Figures 2B-D)** (**52**). This shared strand, which is subtly extended, displays a signature curvature with a kink in the β-strand contributed by a conserved glycine or, occasionally, a proline. This makes this shared strand a “structural signature” in these SBBs, with each half (N and C-termini) forming stabilizing interactions with β-strands of both the orthogonally packed β-sheets, allowing the two sheets to form a semi-open barrel-like structure (**Figures 2B-D**) (**52**). For example, in a typical SH3-like small β-barrel, βS2 is shared between the two sheets (**Figure 2D**). Further signatures of SBB-like domains are typified by β-sheet A, which is sequentially and spatially contiguous, forming a typical β-meander, while β-sheet B is non-contiguous (for example in SH3, β-sheet B is formed by βS1 βS2 (shared) and βS5). The topological connections and structural arrangements of these orthogonally packed sheets serve as the foundation for classifying the SBB group, where each fold exhibits unique topological connections and subtle secondary structure insertions while preserving the broader structural framework of a small β-barrel (**52**).

A comprehensive comparison of NSP7 with established members of the SBB assemblage allowed us to clearly define its relationship with other domains. In NSP7, the three-stranded β-meander is formed by core strands βS3, βS4, and βS5, setting it apart from other SBB members like the OB-fold and the SH3-like fold (**Figures 2C**). The topological connections and orientations of the β-strands differ significantly from the OB and SH3-like folds, and notably, instead of a single β-strand, NSP7 shares two core β-strands, βS4, and βS5, between the two orthogonal sheets (**Figure 2C**). While distinct from these well-known SBB members, the top hits from DALI searches—the HAS barrel domain and the small β-barrel of the 50S ribosomal subunit L21—show a structurally and topologically identical core β-sheet architecture, with two shared β-strands between the orthogonal sheets (**Figure 2C**). In fact, the core of the HAS barrel domain and L21 are structurally superimposable with NSP7, with an average root mean square deviation (RMSD) of less than 4 Å (**Figure 2E**). However, a subtle distinction between the NSP7 and HAS barrel lies in the orientation of βS1 relative to βS2: in NSP7, βS1 runs parallel to βS2, whereas in the HAS barrel, βS1 is arranged antiparallel while the rest of the core barrel is identical (**Figure 2C**). Additionally, these three domains also have distinct and characteristic insert regions and N-terminal/C-terminal extensions (**Figure 2C**). NSP7 features an N-terminal helical bundle, while L21 usually has an N-terminal β-hairpin slightly tilted away from the core sheets and another β-hairpin insert extending from the shared core β-strands βS5 and βS6 of the β-barrel. In contrast, the HAS barrel lacks an N-terminal helical bundle or β-hairpin inserts but usually has an additional strand, βS6, as a C-terminal extension, stacked antiparallel to βS3 through a long loop extending from βS5. The HAS barrel also possesses a helical insert in the same position as the β-hairpin insert between βS5 and βS6 in L21 **(Figure 2B and 2C)**.

Although earlier studies imprecisely classified the L21 ribosomal protein as an SH3-like small β-barrel domain (**53**), our comparative analysis clarifies the structural and topological differences between SH3-like β-barrels and the L21 ribosomal protein, along with its structural homologs— the HAS barrel and arterivirus NSP7. In SH3-like β-barrels, the β-meander is formed by core strands βS2-βS3-βS4, with βS2 shared between the orthogonal sheets (**Figure 2B and 2D**). In contrast, L21 contains a β-meander composed of βS4-βS5-βS6, which structurally aligns with βS3- βS4-βS5 in the β-meander of both arterivirus NSP7 and HAS barrel domains. Our analysis shows that the closest homologs to arterivirus NSP7 are the L21 ribosomal protein and the HAS barrel, although all these domains retain their subtle and unique features (**Figure 2C**). Moreover, the amino acid residues within the NSP7 β-barrel build a hydrophobic core, stabilized by one or two conserved hydrophobic residues from each β-strand, which also serves as a defining characteristic of all SBB domains. Together, we classify arterivirus NSP7 as a definitive member of the broader SBB assemblage, identifying it as an arterivirus-specific (based on its absence in other nidovirales) small β-barrel domain with unique features. Given that NSP7’s closest structural homologs—the HAS barrel and the ribosomal β-barrel L21 – are known to stabilize large protein-protein or protein-nucleic acid complexes (**53–56**), we suggest that NSP7 likely plays a role in viral RNA binding and(or) nucleoprotein complex stabilization, which may be critical to the arterivirus replication cycle. While recent studies lack detailed insights into specific pathways, localization, and interaction mechanisms, a body of earlier works support our inferences, suggesting that NSP7 is essential for interactions with both host and viral proteins and plays a significant role in the overall assembly of the replication-transcription complex (RTC) (**57,58**). With the classification of NSP7 as a SBB domain, focused studies on their modes of action and its potential interaction with the viral RNA and other proteins can be explored.

### Comprehensive characterization of ORF1b – illustrating newly identified and highly derived domains across mammalian and non-mammalian arteriviruses

ORF1b in all previously studied arterivirus genomes is known to encode at least three conserved domains: NiRAN + RdRp, Zinc-binding + Helicase, and NendoU (**12–16**). In mammalian arteriviruses, both the overall length and domain organization of ORF1b are conserved, featuring these three domains followed by the NSP12 domain, which is unique to mammalian arteriviruses (**Figure 1**). Although NSP12 is recognized as encoding a globular domain (**17**), its structural and functional roles are not yet fully understood. Here, we provide evidence that NSP12 is a highly derived version of the MTase-like domain, with N7-MTases from coronaviruses and other nidovirales as its closest homologs. In contrast, non-mammalian arteriviruses show greater variability in the domain composition of ORF1b, including the sporadic presence of two newly identified domains highlighted in this study (see subsequent sections). Additionally, the equivalent of ORF1b is sometimes fused with ORF1a, leading to inconsistencies in nomenclature and the number of ORFs for non-mammalian arteriviruses. To maintain clarity, we designate these equivalent regions as ORF1b and highlight their relationship to mammalian arteriviruses, as illustrated in **Figure 1**.

### Identification of ExoN and canonical 2’-O-MTase in ORF1b of non-mammalian arterivirus

While NSP14 in coronaviruses encodes an ExoN domain essential for proofreading and replication fidelity (**59,60**), earlier studies indicated the absence of this domain in mammalian arteriviruses. Our comparative analysis confirms this finding but uncovers an ExoN domain in five of seven non-mammalian arterivirus genomes (**Figure 1B and 3A**). Structure similarity searches using AF3-predicted arterivirus domains as seeds in DALI identified homologs from coronaviruses (PDB ID – 7EGQ) and various other exonucleases from prokaryotes and eukaryotes. Sequence-structure comparisons with the recovered homologs demonstrated the retention of RNase-H-like fold and conservation of the five catalytic residues (DEDDH) that coordinate Mg²⁺ ions, confirming that ExoN domains in non-mammalian arteriviruses are enzymatically active (**61,62**) **(Figure 3A)**. Notably, unlike coronaviruses, where NSP14 universally includes both the ExoN domain and a guanine N7-MTase, non-mammalian arteriviruses completely lack an N7-MTase. Their ExoN domain is instead located between the helicase and NendoU domains, with no intervening regions that could potentially encode a functional N7-MTase (**Figure 1B**). Interestingly, while mammalian arteriviruses lack an ExoN domain, the NSP12 at the C-terminal end of ORF1b closely resembles the NSP14 N7-MTase of coronaviruses (see next sections). Thus, in contrast to the bifunctional NSP14 ExoN+N7-MTase module found in coronaviruses, the arterivirus genomes present a unique evolutionary scenario, where non-mammalian arteriviruses retain the ExoN domain without an N7-MTase, while mammalian arteriviruses maintain a highly derived N7-MTase-like domain despite lacking an ExoN (discussed in the next section) (**Figure 1B**).

**Figure 3.**
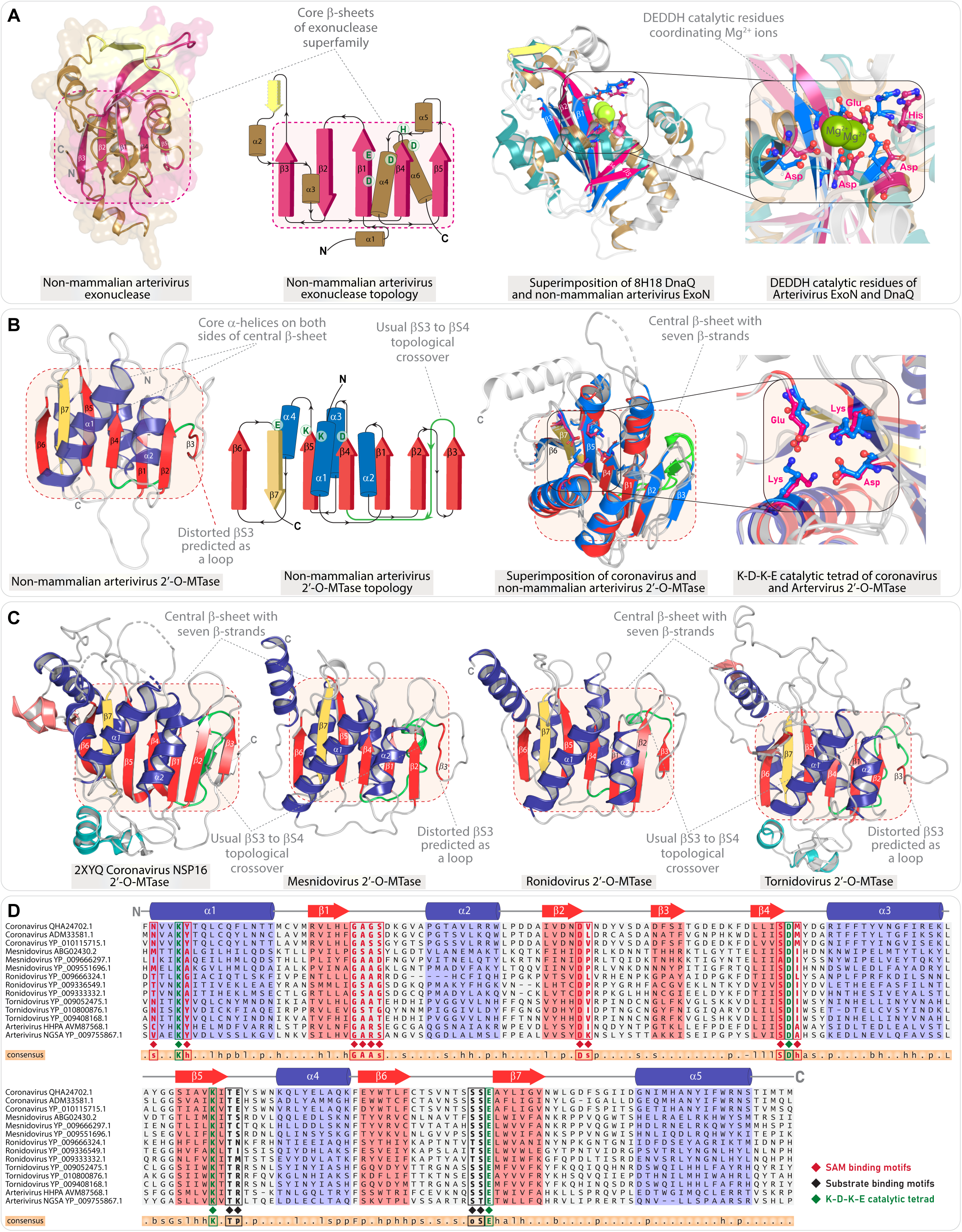
Conserved structural and sequence features of ExoN and 2′O-MTase domains in non-mammalian arteriviruses. **(A)** The first two images in the panel display the representative AF3-predicted 3D structure and the corresponding schematic topology of the exonuclease (ExoN) domain from non-mammalian arteriviruses. The structural superpositions on the right show the alignment of the arterivirus ExoN domain with the DnaQ exonuclease (PDB ID: 8H18), with a zoomed-in view (far right) highlighting the alignment of the DEDDH catalytic residues. **(B)** The first two illustrations in the panel depict the representative AF3-predicted 3D structure and the corresponding schematic topology of the 2ʹO-MTase domain from non-mammalian arteriviruses. The structural superpositions on the right illustrate the alignment of the 2ʹO-MTase domains from arteriviruses and coronaviruses (NSP16 2ʹO-MTase), with a zoomed-in inset (far right) emphasizing the catalytic K-D-K-E tetrad. **(C)** Representative 3D structure of 2’-O-MTases from other Nidovirales. **(D)** Representative multiple sequence alignment of 2ʹO-MTases across all Nidovirales, highlighting key conserved motifs for SAM binding, substrate binding, and the K-D-K-E catalytic tetrad.

While all non-arterivirus Nidovirales members possess at least one 2ʹ-O-Mtase, it remains unclear whether arterivirus genomes encode a bona fide functional MTase domain, as an earlier study identified MTase-like signatures in the HHPA and NGSA genomes (**63**), but did not decidedly confirm their actual presence using comparative sequence-structure analysis. In this study, using AF3 predictions and in-depth comparative analysis, we here unambiguously demonstrate the presence of 2’-O-MTase in three non-mammalian arterivirus genomes (Wuhan Japanese halfbeak arterivirus – WJHA, HHPA and NGSA). Notably, these are retained in only these three non-mammalian arteriviruses and are exclusively absent in the mammalian arteriviruses. These domains interestingly occupy the same position in the genome, C-terminal to the NendoU domain, as the NSP12 in mammalian arteriviruses (**Figure 1B**). From a comparative standpoint, these domains retain the defining features of the NSP16-encoded S-adenosyl methionine (SAM)- dependent 2’-O-MTase found in coronaviruses, including the canonical Rossmann fold and the key residues for binding SAM and m7GpppA (**64**) **(Figure 3B-D)**. Notably, unlike the N7-MTases, the 2’-O-MTases are highly conserved, maintaining most of their characteristic features consistently across all 2’-O-MTases identified in Nidovirales **(Figure 3B-D)**. Structurally, these domains retain the core 2’-O-MTase topology (αH1-βS1-αH2-βS2-βS3-βS4-αH3-βS5-αH4-βS6-βS7-αH5), including N-terminal helices, and inserts following βS3 and βS4. In a comparative context, it is noteworthy that coronaviruses possess both the NSP14 N7-MTase and NSP16 2ʹ-O-MTase, which together form the 5ʹ-end mRNA cap structure essential for evading the host immune system and ensuring the stability of the viral genome within the host (**65–70**). While the retention of both 2ʹ-O-MTase and N7-MTase is not a conserved feature across all Nidovirales, arteriviruses represent a unique case within this order. Canonical 2ʹ-O-MTases are exclusively present in three non-mammalian genomes, while a highly derived version of the N7-MTase-like domain is specific to mammalian arteriviruses.

### Structural and functional insights into NSP12, an inactive N7-MTase-like domain: Conserved core elements and unique variations relative to other Nidovirale N7-MTases

Though this study identifies 2ʹ-O-Mtases in a few non-mammalian arteriviruses, the lack of an MTase domain in mammalian arteriviruses has so far presented an incomplete model of arterivirus biology. Experimental data have detected methylated nucleosides at the 5ʹ end cap of simian hemorrhagic fever virus (SHFV) (**71**), yet prior research has not clearly identified any NSPs with an MTase domain questioning this finding. Adding to this conundrum, a later investigation proposed that mouse lactate dehydrogenase-elevating arterivirus (LDV) might produce uncapped mRNA (**72**), supporting the absence of an MTase domain, leading to inconsistencies in our understanding of arterivirus capping mechanisms. Comparatively, NSP12 located at the 3ʹ end of ORF1b was considered the most likely candidate for a functional MTase domain, as similar genomic positions in other MTase-containing nidovirales are linked to functional MTases. This led to both sequence-based comparative analysis and experimental testing of NSP12 as a possible MTase domain. However, no MTase activity was observed in vitro during assays, even with the addition of other NSPs that could potentially function as cofactors. The authors suggested that either the appropriate cofactors were not included or that NSP12 may require additional cofactors to activate its MTase activity. In addition to these findings, the study showed that suppressing NSP12 expression resulted in a complete absence of viral progeny, underscoring its critical role in arterivirus replication and survival (**17**).

Here, using AF3-predicted structures of all 23 mammalian arteriviruses and detailed structural similarity analyses with DALI and HHPRED, we demonstrate that NSP12 consistently exhibits a conserved core six-stranded β-sheet, forming a Rossmann fold typical of an MTase-like domain, but with subtle yet important deviations. (**Figure 4A**). In typical Rossmann fold MTases, the central β-sheet comprises seven β-strands arranged in the sequence: 6↑-7↓-5↑-4↑-1↑-2↑-3↑. Key features of this central β-sheet include: (i) a characteristic topological crossover where βS3 at the rightmost end leads to βS4 in the centre; (ii) a β-hairpin at the C-terminal end, formed by βS6 and βS7, with βS7 inserted anti-parallel between βS5 and βS6, which serves as a key synapomorphy of Rossmann fold MTases. (**73–78**). In NSP12, the core sheet consists of six strands (5↑-6↓-4↑-3↑-1↑-2↑), with the loss of the typical βS3 causing βS2 to become the rightmost strand, and the topological crossover occurs from βS2 to βS3 at the centre (**Figure 4A**). Aside the absence of βS3, the core sheet of NSP12 precisely aligns with the central β-sheet architecture of canonical MTases retaining all other structural synapomorphies including the C-terminal hairpin. Besides the core region, NSP12 contains a C-terminal α-helix positioned parallel to the core sheet, followed by an extended loop that leads to a Zn finger module stacked in front of the core sheet (**Figure 4A**). In some instances, a small β-strand insert is observed between αH1 and βS1, which is stacked together with the β-hairpin of the Zn finger module, forming a three-stranded sheet that ends with a C-terminal helix (**Figure 4A**).

**Figure 4.**
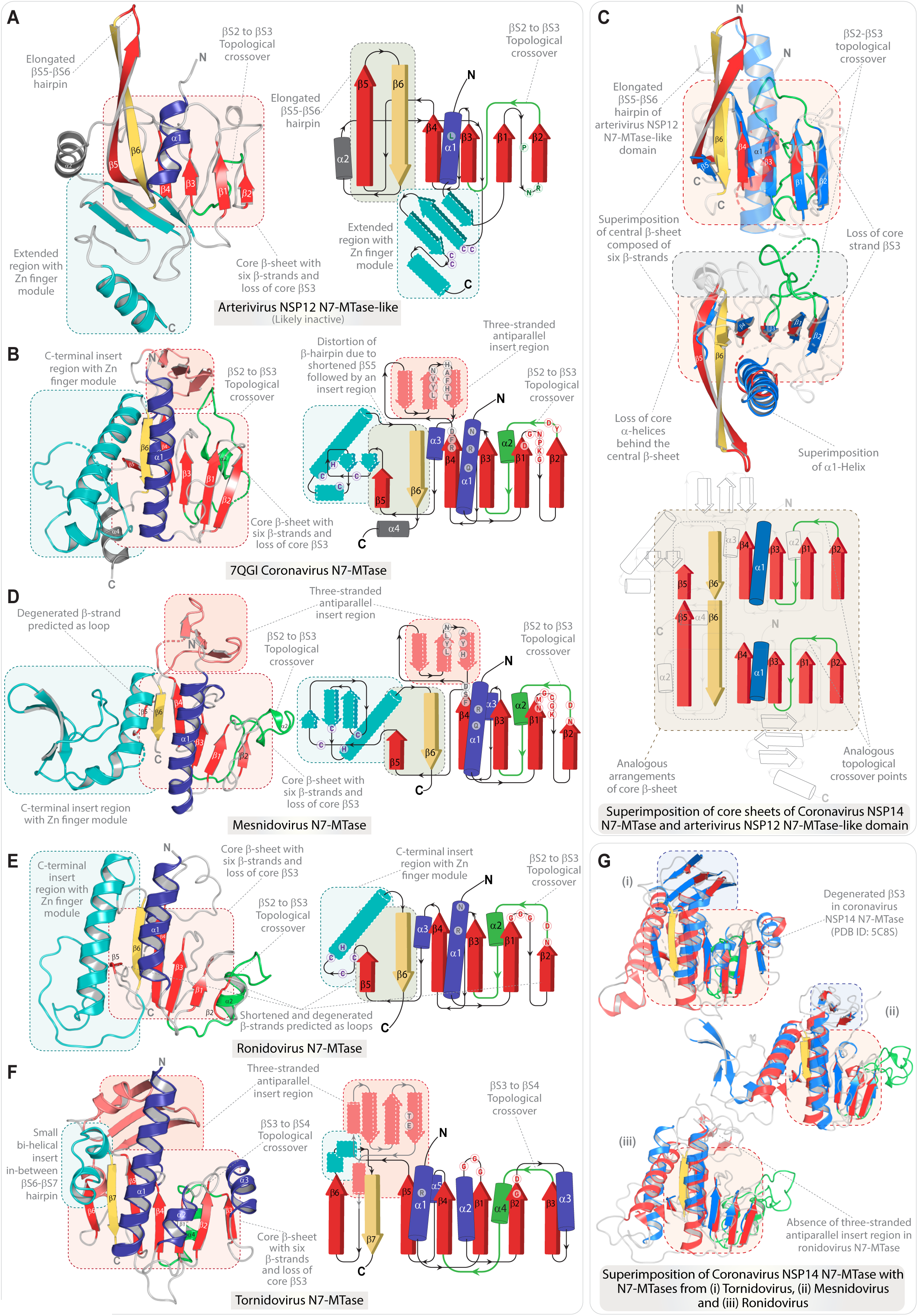
Structural features of the arterivirus NSP12 N7-MTase-like domain and its comparison with Nidovirale N7-MTases. Panels **(A)** and **(B)** display the 3D structure and corresponding topology of the arterivirus NSP12 N7-MTase-like domain (AF3 predicted) and the coronavirus NSP14 N7-MTases (PDB ID: 7QGI). **(C)** The illustration on the top shows the structural superposition of the arterivirus NSP12 N7-MTase-like domain with the coronavirus NSP14 N7-MTase (side view), while the middle unit presents a top view, focusing primarily on the superposition of the core sheet and the anterior core αH1. The illustration at the bottom of the panel shows the topological comparison of the core β-sheet of the arterivirus NSP12 N7-MTase-like domain with the coronavirus NSP14 N7-MTase. The shared core sheet and the αH1 are colored. **(D-F)** 3D structure and corresponding topology diagrams of N7-MTases from mesnidovirus, ronidovirus, and tornidovirus. **(G)** Structural superposition of coronavirus NSP14 N7-MTase with N7-MTases from (i) tornidovirus, (ii) mesnidovirus, and (iii) ronidovirus.

We conducted an extensive survey and comparisons to determine whether similar domains are present in other nidovirales. Our structural and sequence analyses revealed several notable findings:

i. At the structural level, we found that both the arterivirus NSP12 and coronavirus NSP14 N7-MTase domains exhibit the same structural framework, which can be summarized as a derived Rossmann fold. This is specifically defined by the absence of the core βS3 strand, which leads to a βS2-to-βS3 crossover. Consequently, the core β-sheet consists of four parallel β-strands, followed by the C-terminal β-hairpin (**79–84**) (**Figure 4A-C)**. In some experimentally determined structures (PDB IDs: 5C8S and 7N0D), the core β-sheet includes seven β-strands, however βS3 is small and degenerated, containing only two residues. While this forms the typical βS3-βS4 topological crossover, βS3 is not consistently resolved in all structures, with sequence alignment supporting its degeneration (**Figure 5**). Additionally, both domains show a reduction in core helices anterior to the core sheet, retaining only αH1, deviating from the typical and packed Rossmann fold αβα sandwich structure. Behind the sheet, arterivirus NSP12 lacks helices, while coronavirus NSP14 retains two short helices (αH2 and αH3). Building on these observations, we found that N7-MTases from mesnidoviruses and ronidoviruses share identical core topology features, including a six-stranded architecture, the absence of βS3, and a reduction in core helices, with minimal deviations **(See Figure 4D and 4E and Table 1)**.
ii. Among all nidovirale N7-MTases, only the Tornidovirineae N7-MTase retains most of the canonical features of Rossmann fold MTases (**63,85**). This includes the retention of the βS3 strand, the characteristic βS3-to-βS4 crossover, and the typical arrangement of five parallel strands followed by a C-terminal β-hairpin (**Figure 4F and Table 1**). Additionally, it preserves three helices in front of the core sheet and two short helices at the rear, making its structure the closest to canonical Rossmann fold MTases (**Figure 4F)**.
iii. In addition to these features, the N7-MTases of nidovirales are marked by two key regions that are positionally conserved yet only sporadically retained across the group (**Figure 4A-G)**: (i) a Zn finger module at the C-terminal end, and (ii) a three-stranded antiparallel insert region that extends outward above the core β-sheet from βS4 (βS5 in Tornidovirineae), forming a cap-like structure and a small pocket region. Previous studies have described this as a “hinge” domain located at the interaction interface between the N7-MTase and ExoN domains within the N7-MTase+ExoN heterodimer in coronavirus, contributing additional residues for substrate binding (**79–81,86**). However, these insert regions are not consistently present across nidovirales. The Zn finger module is found in arteriviruses (though not fully conserved), coronaviruses, mesnidoviruses and ronidoviruses, while the three-stranded antiparallel β-sheet insert is present in all nidovirales except for arteriviruses and ronidoviruses (**Figure 4A-G**). Thus, the potential role of the Zn finger module in RNA binding or protein-protein interactions likely varies across virus clades, suggesting a context-dependent, clade-specific role that may be dispensable in certain groups. Similarly, the three-stranded antiparallel β-sheet does not represent a conserved mechanism across all Nidovirales.
iv. Each of these domains also exhibits subtle, clade-specific features **(Figure 4 and Table 1)**. For example, in NSP12, the βS5-βS6 hairpin is notably longer, spanning 12-13 amino acids, while in the coronavirus N7-MTase, βS5 is shorter, followed by an insertion leading to an antiparallel βS6, typically six residues long (**Figure 4A and 4B**). In mesnidoviruses and ronidoviruses N7-MTases, βS5 is shorter or often predicted as a loop in AF3 models. Additionally, the Zn finger module at the C-terminal region exhibits clade-specific variations, making it not fully superimposable. Notably, arteriviruses feature Zn2+ ion chelation by C4-type zinc fingers, whereas coronaviruses, mesnidoviruses, and ronidoviruses display C3H-type zinc fingers (**Figures 4 and 5 and Table 1**).

**Figure 5.**
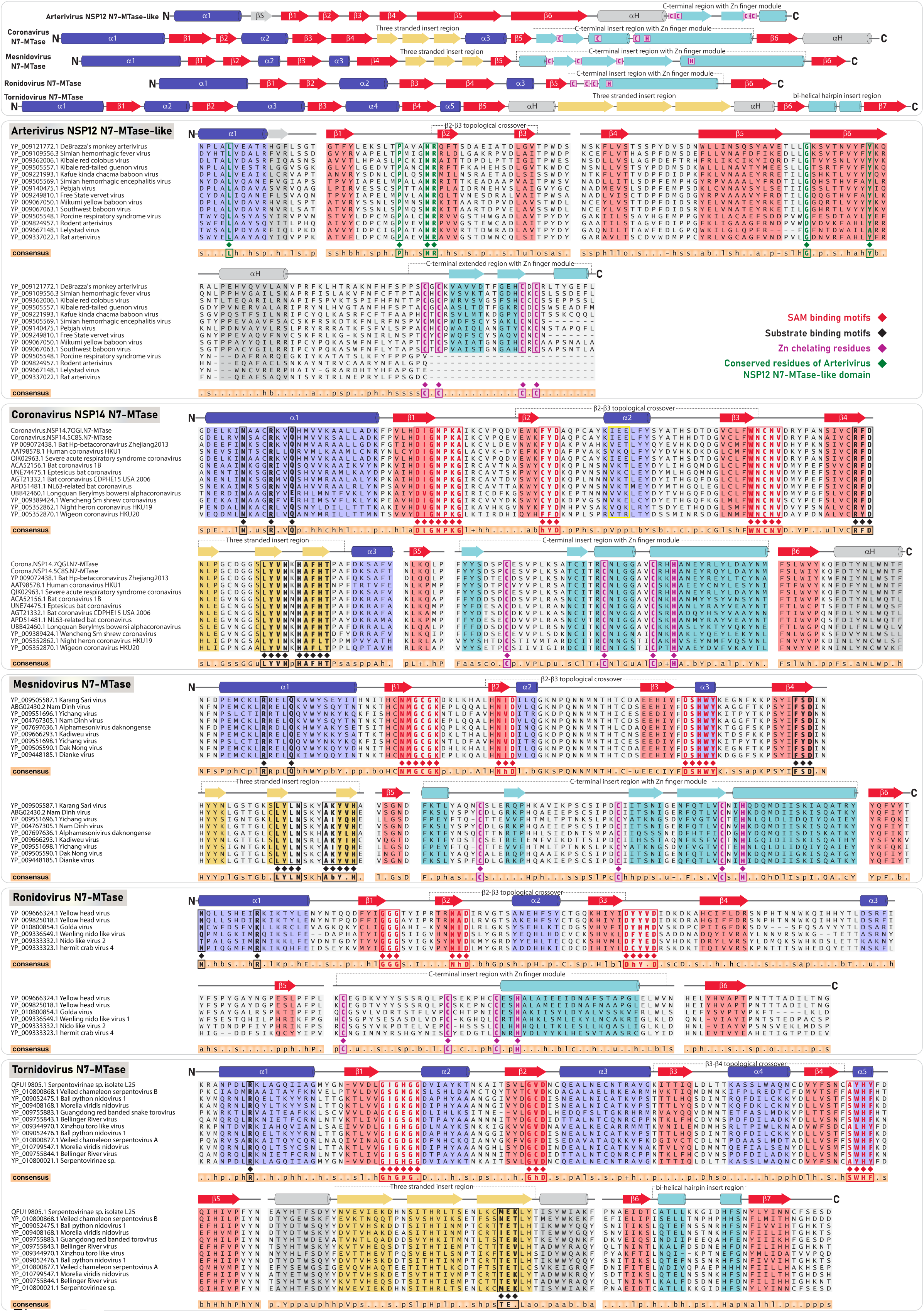
Multiple Sequence Alignment of the arterivirus NSP12 N7-MTase-like domain with N7-MTases across Nidovirales. The top panel presents a linear topological comparison of the arterivirus NSP12 N7-MTase-like domain with N7-MTases across Nidovirales. The core α-helices and β-sheet forming the Rossmann fold are shown in blue and red, respectively, with the three-stranded insert in yellow and the C-terminal insert in teal. Secondary structural elements positioned outside of the αβα sandwich is colored grey. The panels below display representative multiple sequence alignments of N7-MTase domains from all five distinct Nidovirales families, with secondary structural elements colored as described. In the coronavirus N7-MTase, a short, inconsistently predicted βS3, which occasionally causes the structure to resemble and portray a seven-strand sheet, is highlighted with a yellow border. A consensus sequence is indicated at the bottom of each alignment. SAM-binding motifs, substrate-binding motifs, and Zn-chelating residues are highlighted in red, black, and magenta, respectively. Excluding the Zn-chelating residues, the SAM and substrate-binding (guanine-binding) residues exhibit erosion and lack conservation in arteriviruses. Arterivirus-specific conserved residues are highlighted in green.

**Table 1.** Synapomorphies and distinctive variations of N7-MTases in Nidovirales.

|  | **Tornidovirineae**<br>**N7-MTase** | **Cornidovirineae**<br>**N7-MTase** | **Mesnidovirineae**<br>**N7-MTase** | **Ronidovirineae**<br>**N7-MTase** | **Arteriviridae**<br>**NSP12 N7-MTase-**<br>**like** |
| --- | --- | --- | --- | --- | --- |
| **Core topology excluding inserts** | αH1-βS1-αH2-βS2-αH3-βS3-(αH4)-βS4-(αH5)-βS5-βS6-βS7. αH4 and αH5 are shortened. | αH1-βS1-βS2-(αH2)-βS3-βS4-(αH3)-βS5-βS6. αH3 is shortened. | αH1-βS1-βS2-(αH2)-βS3-(αH3)-βS4-βS5-βS6. αH2 and αH3 are shortened. | αH1-βS1-βS2-(αH2)-βS3-βS4-(αH3)-βS5-βS6. αH2 and αH3 are shortened. | αH1-βS1-βS2-βS3-βS4-βS5-βS6. |
| **Genomic location** | ORF1a (**63,85**). | NSP14 of ORF1b along with an ExoN, followed by NSP15 NendoU and NSP16 2'-O-MTase domains (**79-81**). | C-terminal end of ORF1b, preceded by an ExoN and followed by 2'-O-MTase (**7**). | C-terminal end of ORF1b, preceded by an ExoN and followed by 2'-O-MTase (**8**). | NSP12 located at the C-terminal end of ORF1b, preceded by a NSP11 NendoU domain (**1**). |
| **Core helices of N7-MTases** | Three core helices αH1, αH2 and αH3 on the anterior side of central β-sheet; Shortened αH4 and αH5 on the posterior side of central β-sheet. | Only one helix (αH1) on the anterior side of central β-sheet; Lacks other helices on the anterior side; Two short helices αH2 and αH3 on the posterior side of central β-sheet (**79-81**). | Only one helix (αH1) on the anterior side of central β-sheet; Lacks other helices on the anterior side; Two short helices αH2 and αH3 on the posterior side of central β-sheet. | Only one helix (αH1) on the anterior side of central β-sheet; Lacks other helices on the anterior side; Two short helices αH2 and αH3 on the posterior side of central β-sheet. | Only one helix (αH1) on the anterior side of central β-sheet; Lacks other helices on the anterior side; Loss or short helices on the posterior side of central β-sheet. |
| **Central β-sheet architecture** | Canonical seven-stranded core β-sheet with a typical Rossmann fold as observed in MTases (**63,85**); Usual βS3-βS4 topological crossover. | Six-stranded core β-sheet architecture; βS2-βS3 topological crossover due to degeneration of typical βS3; Degenerated core βS3 and shortened βS5; βS3 is occasionally represented by a small strand in PDB ID 5C8S and 7N0D (**79,82**), forming a seven β-stranded central sheet (not consistent) | Six-stranded core β-sheet architecture; βS2-βS3 topological crossover due to degeneration of core βS3; Shortened βS5 is predicted as a loop in a few AF2 structures. | Six-stranded core β-sheet architecture; βS2-βS3 topological crossover due to degeneration of core βS3; Shortened βS5 is predicted as a loop in a few AF2 structures. | Six-stranded core β-sheet architecture; βS2-βS3 topological crossover due to degeneration of core βS3. Instead of shortened βS5, the βS5 is elongated forming an elongated hairpin with βS6. |
| **Three-stranded anti-parallel β-sheet insert region** | Three stranded antiparallel β-sheet insert region after core βS5; includes additional small helices flanking the sheet. | Three stranded antiparallel β-sheet insert region after core βS4 (**79-81**). | Three stranded antiparallel β-sheet insert region after core βS4. | Absence of insert region. βS4 directly leads to a shortened αH3 and βS5 with no β-sheet insert region. | Absence of insert region. βS4 directly leads to βS5-βS6 hairpin with no insert region. |
| **C-terminal insert region with Zn finger module** | Small bi-helical hairpin insert region between core βS6 and βS7. Absence of Zn finger module. | Insert region with two small β-strands and two helices between core βS5 and βS6; Contains a conserved | Insert region with three antiparallel β-sheet followed by a helix: Insert occurs between core βS5 and βS6; Contains a | Insert region with one helix between core βS5 and βS6; Yet the loop region connecting the shortened βS5 and | Elongated βS5-βS6 hairpin with no insert region; The Zn finger module occurs as a C-terminal extension |
| | | Zn finger module (C3H type). The insert after βS5 subtly distorts the typical C-terminal β-hairpin of Rossmann fold MTases. Short helix after βS6 | conserved Zn finger module. The insert after βS5 subtly distorts the typical C-terminal β-hairpin of Rossmann fold MTases. | insert helix encodes the C3H type Zn finger module. Shortened βS5 distorts the typical C-terminal β-hairpin of Rossmann fold MTases. | (C4 type). An additional strand as an insert between αH1 and βS1 is stacked against the strands of the C-terminal extension. |
| **Key SAM binding regions** | Glycine rich loop with a characteristic motif GxGxG, on top of βS1. Conserved D in motif G[h]D, on top of βS2 (h: hydrophobic residue). Conserved motif WH[Y/F], on top of βS4 (**63,85**). | Glycine rich loop with a subtly distinct motif: DI followed by GNPKG, on top of βS1; Conserved D in motif [F/C]YD, on top of βS2; Conserved motif WNCNV on top of βS3 (**79-82**). | Glycine rich loop with a subtly distinct motif NMGCGK, on top of βS1; Conserved D in motif N[I/V]D, on top of βS2; Conserved motif DSHWY on top of βS3. | Glycine rich loop with a distinct motif GGG, on top of βS1; Conserved D in motif N[h]D, on top of βS2 (h: hydrophobic residue); Conserved motif D[h]Y[x]D on top of βS3. | No conservation of residues in structurally superimposable region. |
| **Key substrate binding regions** | Conserved R in αH1. Conserved motif TE[T\K], in the three stranded β-sheet insert region (**63,85**). | Conserved N, R and [Q/E] in αH1; Conserved motif R[F/Y]D on top of βS4; LY[V/I]N and HAF[H/L]T in the three stranded β-sheet insert region (**79-81**). | Conserved R and Q in αH1; Conserved motif F[S/Y]D on top of βS4; LYLN and A[b]Y[x]H motifs in the structurally superimposable three stranded β-sheet insert region. "b" denotes basic, "x" any amino acid. | Partially conserved N and a conserved R in αH1; Loss of conserved motif on top of βS4 and loss of three stranded β-sheet insert region. | No conservation of residues in structurally superimposable region. The only conserved residues include a L αH1, P and a "NR" dyad in the loop connecting βS1-βS2. |
| **Additional comments and remarks** | Only N7-MTase domain within Nidovirales that exhibit seven-stranded Rossmann fold with βS3-βS4 crossover; However, shares the three stranded β-sheet insert region and multiple key binding sites with derived N7-MTases of other nidovirales. | This derived six-stranded version is retained in all Nidovirales except for Tornidovirus which exhibits a more canonical version. However, the domain retains catalytic features. Shown to be an active MTase domain (**79-81**). | This derived six-stranded version is retained in all Nidovirales except for Tornidovirus which exhibits a more canonical version. However, the domain retains catalytic features. | This derived six-stranded version is retained in all Nidovirales except for Tornidovirus which exhibits a more canonical version. However, the domain retains catalytic features. | While retaining the derived six-stranded version, the domain exhibits further variations with an unusually extended C-terminal β-hairpin and no conservation of catalytic residues. Likely inactive domain. |

At the sequence level, all Nidovirales N7-MTases, except for arterivirus NSP12, retain the key catalytic residues for SAM and substrate binding **(Figure 5 and Table 1)** (**64,79,81,87**), though some require additional NSPs as co-factors for MTase activity (**79,80,88–90**). In contrast, NSP12 from arterivirus lacks sequence conservation in the SAM-binding and substrate-binding sites, marking it as a unique, inactive MTase-like domain specific to mammalian arteriviruses (**Figure 5**). While structural comparisons suggest a potential role in mRNA capping like other N7-MTases, the absence of conserved and key catalytic residues indicates no enzymatic activity, and the key regions exhibit no significant selective pressure to retain these residues (**Figure 5**). Interestingly, the only conserved residues are a lysine located in the middle of αH1, a proline found in the center of the loop leading to βS2, and an intriguing asparagine-arginine dyad at the end of the loop just before the start of βS2 (**Figure 5**). However, these residues are positioned away from the typical SAM and substrate-binding sites.

Together, we propose that NSP12 from arteriviruses encodes a derived and inactive N7-MTase-like Rossmann domain. While this inactive MTase domain seems unique to arteriviruses within nidovirales, similar domains are not uncommon, as seen with the inactive MTase domain at the C-terminal end of NSP2 in alphaviruses (**91–93**) **(Supplementary Figure S3)**. Although catalytically inactive due to the loss of SAM-binding motifs, the alphavirus inactive MTase domain retains the canonical Rossmann fold, potentially forming an RNA-binding interface with the N-terminal RNA helicase and protease domains of NSP2, and the downstream NSP3, which includes an ADP-ribose binding macro-domain and a Zinc-binding domain (**76,93–95**). While NSP2, comprising an RNA helicase, protease, and iMTase domain, is known to degrade the RNA polymerase II subunit RPB1, inhibiting host transcription and reducing IFN-stimulated gene (ISG) expression (**96**), recent studies also underscore the critical role of its C-terminal inactive MTase domain in host immune evasion. This domain has previously been shown to selectively inhibit the JAK/STAT signaling pathway, facilitating immune evasion by promoting CRM-1 (chromosome region maintenance 1 export receptor)-mediated nuclear export of the STAT1 transcription factor, thereby preventing the induction of ISG expression (**97,98**). Remarkably, the iMTase domain maintains its inhibitory function independently of the other NSP2 domains, underscoring its specific role in alphaviruses (**97**). Earlier studies also suggest that the iMTase may function as a dominant-negative, hindering STAT1 methylation by disrupting host MTases (SETD2), which are essential for STAT1 phosphorylation and activity (**97,99**). While the precise mechanisms and roles remain to be fully explored, it is evident that the domain is recruited and maintained for immune evasion in alphaviruses. Likewise, these observations suggest that the retention of the inactive MTase domain with its preserved Rossmann fold in arterivirus NSP12 may enable a specialized function in immune evasion by modulating the host immune response, or it may simply aid in RNA binding within the viral replication machinery.

### Divergence of structural proteins in arteriviruses: host-specific evolution and variability across mammalian and non-mammalian genomes

Though the C-terminal genomic organization and the number of ORFs are largely conserved among mammalian arteriviruses, our analysis reveals that the structural proteins they encode— particularly the envelope glycoproteins (GPs) and membrane (M) proteins—exhibit signatures of rapid evolution and considerable variability in sequence and structure. This trend also applies to structural proteins from non-mammalian arteriviruses, which demonstrate significant sequence diversity across different host species. Indeed, HHPRED searches using these sequences and DALI searches utilizing AF3-predicted non-mammalian structural proteins found no reliable hits from other viruses. By analyzing datasets that include structural proteins from both mammalian and non-mammalian arteriviruses, we observed a strong pattern of host-specific divergence through CLANS-based clustering and phylogenetic analysis (**Figure 6**). In mammalian arteriviruses, this divergence is particularly evident between primate and non-primate hosts, with structural proteins from these groups forming well-defined and distinct clusters. Phylogenetic trees for each of the analyzed structural proteins consistently validate this observation (**Figure 6)**. Furthermore, studies have demonstrated that primate arteriviruses possess two copies each of GP2, GP3, and GP4, resulting from a duplication event, and the tree reconstruction shows that the ancestral and duplicate copies form distinct clusters (**20,100,101**). The observed differences in sequence and AF3 predicted structures underscore previous findings that the ancestral and duplicated glycoproteins, unique to primate arteriviruses, fulfill independent and non-redundant functions (**100**).

**Figure 6.**
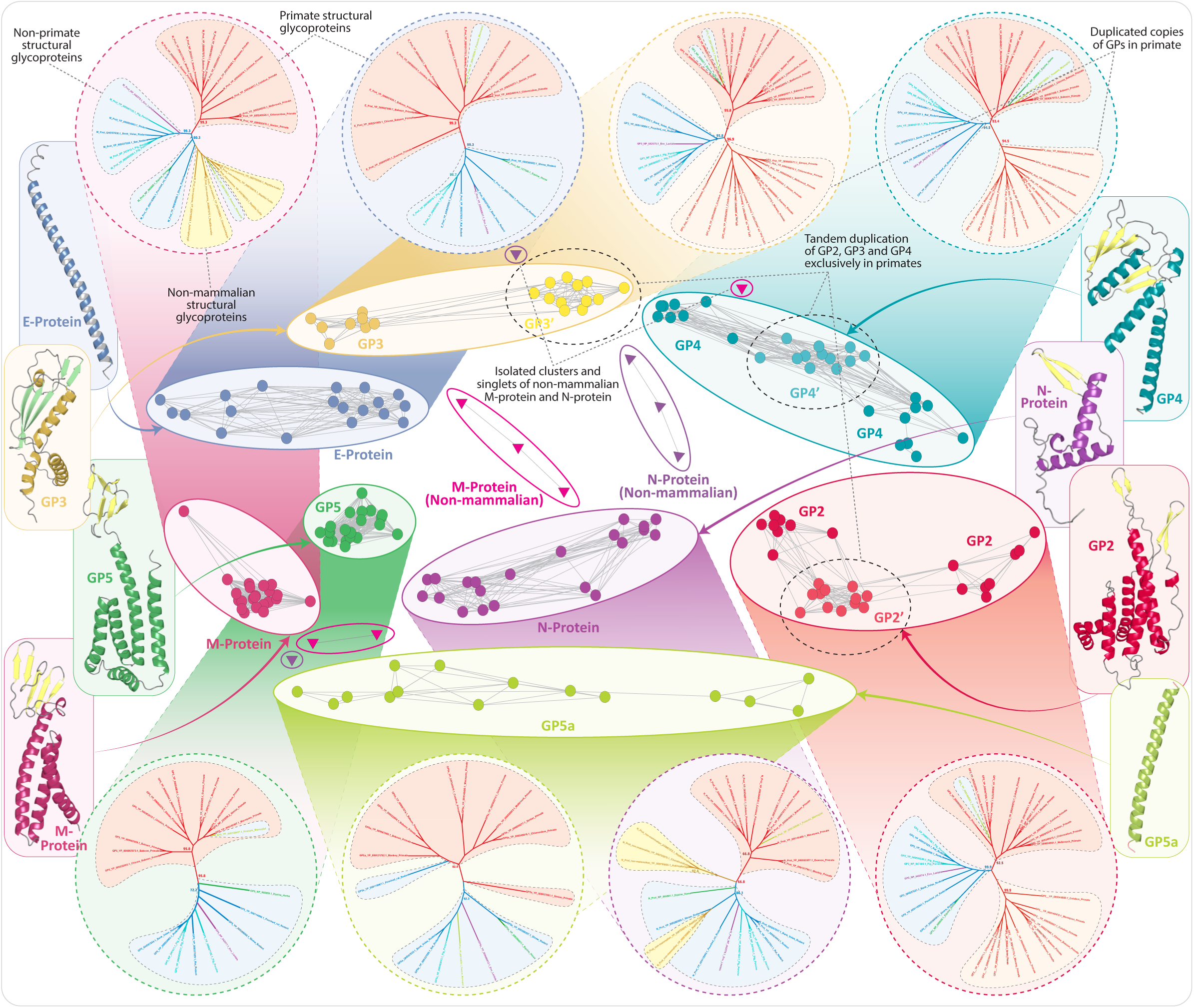
CLANS-based clustering and phylogenetic analysis of arterivirus structural glycoproteins. The central section of the figure displays the overall CLANS-based clustering patterns of structural glycoproteins from both mammalian and non-mammalian arteriviruses. Using CLANS, all-versus-all pairwise sequence similarity-based clustering was conducted, with each glycoprotein cluster shown in distinct colors. Alongside each CLANS cluster, the corresponding maximum likelihood (ML) trees illustrate evolutionary relationships. The ML trees show clear clustering patterns that reflect host-specific divergence, especially between primate and non-primate hosts—primate clusters are highlighted in red, non-primate clusters in blue. Likewise, the non-mammalian GPs are distinct, and their clusters are highlighted in yellow. The ML trees are presented within dotted circles, with background color matching their respective CLANS clusters. Similarly, the structures of each glycoprotein are displayed in solid boxes, with colors consistent with their respective CLANS clusters.

Unlike mammalian arteriviruses, non-mammalian arteriviruses show no conservation in either genomic organization or the number of ORFs encoding structural proteins. While their genome sizes generally range from 13 to 15 kb, similar to their mammalian counterparts, some non-mammalian arterivirus genomes extend beyond this, reaching up to 18 kb (**Figure 1B**). Notably, these include the arteriviruses from ray-finned fish – WJHA, Guangdong greater green snake arterivirus (GGGSA), and HHPA. Despite their large size, these viruses have fewer ORFs than mammalian arteriviruses. This reduction in ORF count is partly due to ORF fusion, such as the frequent merging of ORF1b with ORF1a, which leads to a decrease in the overall number of ORFs (**Figure 1B**). We also predict that similar fusions have likely occurred in ORFs encoding structural proteins, as some ORFs are unusually long compared to those found in mammalian arteriviruses (**Figure 1B**). For instance, certain GP-encoding ORFs from GGGSA, HHPA, Chinese broad-headed pond turtle arterivirus – CBPTA, and TSHSA are significantly larger, ranging between 568 and 799 amino acids. AF3 predictions suggest that these ORFs contain at least one well-defined GP-like domain, along with additional segments that may encode a second GP-like domain (**Supplementary Figure S4**). However, many of these segments are too disordered to accurately define their domain compositions. Moreover, five other GP-encoding ORFs from these viruses exceed 300 amino acids, which is in sharp contrast to mammalian arteriviruses, where the longest structural GPs (typically GP2 and GP5) range between 200 and 300 amino acids (**Figure 1B and Supplementary Figure S4**)

Despite their diversity, we were able to classify the majority of these proteins. For instance, all non-mammalian arterivirus genomes, with the exception of WJHA, contain at least two ORFs encoding GP-like structures that maintain key features, such as an extracellular ectodomain, one or more transmembrane segments, and an extended intracellular endodomain loop (**102,103**) (**Supplementary Figure S4**). Through sequence alignments and structural topology analysis, we identified M proteins in at least six non-mammalian arteriviruses (**Figure 1B and Figure 6**). Similarly, the N protein features a disordered, positively charged RNA-binding region at the N-terminus and a C-terminal domain comprised of α-helices and β-strands forming a four-stranded antiparallel β-sheet (**104–106**). Anchoring on these features, we identified N proteins in at least five non-mammalian arteriviruses. Both M and N proteins in these viruses have experienced significant divergence, leading to small, isolated clusters or singlets in CLANS network analysis (**Figure 6**). Phylogenetically, non-mammalian M and N proteins are distinct from their mammalian counterparts, with a few exceptions. While we could not find ORFs encoding structural homologs of M proteins in GGGSA isolates or N proteins in two genomes (HOFA and TSHSA), we advise caution in concluding that these ORFs have been lost, as this may be due to poor genome assemblies. This is further illustrated by the TSHSA genome, which deviates from the typical genomic architecture and includes two atypical ORFs (**22**) located before the canonical ORF1a, encoding a transmembrane segment and PLP2 (**Figure 1B**). Along with the significantly larger GP-like ORFs at its C-terminus, these features may represent artifacts resulting from suboptimal genome assemblies. Additionally, there are eight other structural proteins, ranging from 35 to 205 residues. AF3 predictions suggest these are mostly disordered or consist of helical segments, lacking clear structural homology for precise annotation. While these may represent GPs, we provisionally label them with a question mark as ‘GP?’ (**Figure 1B**). Nevertheless, the overall comparisons of the structural proteins from arteriviruses highlight the dynamic evolution of these viruses and emphasize the influence of host-specific factors in shaping the divergence of structural proteins across different host species (**103,107,108**).

### Overall Functional and Evolutionary Implications

The comprehensive comparative analysis of arterivirus genomes, including seven non-mammalian genomes, alongside the identification of domains unique to mammalian and non-mammalian arteriviruses, provides valuable insights into the evolutionary dynamics and adaptive strategies of these viruses, highlighting the complex landscape of viral replication, immune evasion, and lineage-specific adaptations.

Firstly, the presence of multiple PLP paralogs, such as PLP1α, PLP1β, and primate-specific PLP1γ, alongside the conserved PLP2 with its proteolytic and deubiquitinase activities, underscores the key roles and adaptive evolution of PLPs in mammalian arteriviruses for polyprotein processing and immune evasion (**9–11,40,41**). In contrast, the reduced number of PLPs in non-mammalian arteriviruses—particularly the absence of PLP1β and PLP1γ—indicates a different evolutionary trajectory linked to reptilian or chondrichthyan hosts, with only PLP2 being conserved. Structurally, non-mammalian PLPs, especially PLP2, exhibit subtle deviations that may reflect optimization for their specific hosts. While the deubiquitinase activity of these PLP2s remains to be experimentally validated, their conservation suggests retained functions, including DUB activity. The absence of PLP2 in certain lineages, like the Nanhai ghost shark arterivirus and Hainan Hebius popei arterivirus, may be attributed to suboptimal genome assemblies. Furthermore, the identification of newly characterized domains in ORF1a, particularly globular domains with an α+β topology, points to unique structural and functional innovations. Their lack of homology with known proteins suggests lineage-specific roles, potentially linked to replication, immune modulation, or other viral processes that remain to be elucidated. It is also plausible that some of these domains may perform functions similar to PLPs, warranting further experimental investigation. Additionally, the presence of disordered intervening regions (DIVs) in ORF1a enhances evolutionary plasticity, potentially acting as flexible scaffolds or interaction hubs that enable rapid adaptation to host environments. This structural flexibility may also support the integration of new protein-protein interactions, crucial for assembling replication complexes and evading host immune defenses.

Likewise, the identification of NSP3C as a conserved wHTH domain and NSP7 as a small β-barrel domain in arteriviruses reveals important insights into their likely roles in viral replication processes. The structural resemblance between NSP3C and coronavirus NSP4C suggests a shared function in RNA binding within replication-transcription complexes (RTCs), likely facilitating interactions with single-stranded RNA and supporting the formation of double-membrane vesicles (DMVs) crucial for viral genome replication (**46,48**). This conservation across mammalian and non-mammalian arteriviruses underscores their essential role in the replication process. Meanwhile, the NSP7, part of the small β-barrel (SBB) domain family, shares structural homology with the HAS barrel and L21 ribosomal protein, both known for binding nucleic acids and stabilizing protein-nucleic acid complexes (**53–56**). Its conservation across different hosts highlights its importance in RNA binding or nucleoprotein complex stabilization, critical for arterivirus replication. The unique recruitment of the SBB domain in arteriviruses, which is likely absent (not exclusively identified so far) in other nidovirales, suggests a potentially important and clade-specific role in the replication machinery of arteriviruses. Together, NSP3C and NSP7 are likely essential components of arterivirus replication (especially mammalian arteriviruses), with possible co-evolutionary implications for host-virus interactions and viral persistence, necessitating further investigation into their functions in RNA binding and immune evasion.

The evolutionary divergence of ORF1b between mammalian and non-mammalian arteriviruses highlights distinct replication and immune evasion strategies shaped by lineage-specific adaptations. Mammalian arteriviruses appear to have lost both the ExoN and 2ʹ-O-MTase domains, with the NSP12 MTase-like domain repurposed for a unique function due to its likely loss of enzymatic activity. In contrast, other nidovirales, such as coronaviruses, employ both the N7-MTase + ExoN heterodimer and 2ʹ-O-MTase domains in their mRNA capping mechanisms, indicating that mammalian arteriviruses may not thoroughly rely on mRNA capping for immune evasion. Meanwhile, non-mammalian arteriviruses sporadically retain functional methylase domains. While the N7-MTase domain is absent, the ExoN domain—typically fused with N7-MTases in coronaviruses—is occasionally retained. In these cases, arteriviruses maintain a 2ʹ-O-MTase domain that may compensate for the absence of N7-MTase. Overall, there is no strong conservation of methylase domains across arteriviruses. Notably, among all nidovirales, arteriviruses demonstrate a distinctive and remarkable pattern, largely lacking methylases, with only sporadic instances of 2ʹ-O-MTase present in just three analyzed genomes. A similar lack of conservation of N7-MTases is observed in the Tobaniviridae family, where N7-MTase activity is restricted to non-mammalian isolates (**63**). However, 2ʹ-O-MTases are nearly universally present across Nidovirales, excluding arteriviruses. This diversity and specific retention of these domains suggest that ancestral arteriviruses, or the clade leading to the emergence of arteriviruses, likely possessed them, with mammalian arteriviruses losing both ExoN and the enzymatic function of NSP12, while non-mammalian arteriviruses sporadically retain 2ʹ-O-MTase and ExoN. Nevertheless, the conservation of NSP12 in mammalian arteriviruses, despite its inactivity, indicates its essential role as a replication co-factor rather than as a methylase.

These patterns in phyletic distribution underscore a distinct evolutionary trajectory for 2ʹ-O-MTase and N7-MTase, with the former being more conserved and the latter exhibiting signs of adaptive evolution, likely reflecting involvement in an arms race with the host immune system. Our survey and comparisons suggest that the derived version of the N7-MTase domain was potentially present early in the evolution of Nidovirales, alongside the Zn finger module and the characteristic three-stranded antiparallel β-sheet insert, as well as the loss of core helices. The sporadic retention of these features may indicate that the Zn finger module and antiparallel β-sheet insert were likely lost from the ancestral form in specific nidovirale clades, rather than acquiring them independently. While it remains unclear whether the common ancestor of all Nidovirales N7-MTases possessed the βS3 strand, the loss of βS3 in all but Tornidovirineae suggests that this loss occurred much earlier, prior to the emergence of distinct Nidovirales clades. Tornidovirineae likely retained the version with βS3, whereas other clades preserved derived versions lacking it. Considering the unique evolutionary trajectories of various Nidovirales clades and their diverse host ranges, the specific retention of these characteristic structural features in N7-MTases suggests a highly selective functional role from a structural perspective, which has yet to be fully elucidated. Overall, these findings underscore the significance of domain co-option in the evolution of arteriviruses and the impact of host-virus co-evolution on shaping the diversity of arterivirus proteins between mammalian and non-mammalian arteriviruses.

## Conclusions

The findings presented here, derived from comparative genomics and detailed sequence-structure analysis, elucidate several previously overlooked and unannotated domains in arteriviruses, providing a broader perspective on their evolution. By incorporating all sequenced non-mammalian genomes and reconstructing their protein domain architectures, we performed a comprehensive comparative analysis that highlighted significant differences in functional domain composition between mammalian and non-mammalian arteriviruses, including domains unique to each group. We offer the first classification and structural insights into the NSP3C wHTH and the small β-barrel domain of NSP7 in mammalian arteriviruses. Intriguingly, we identify NSP12 in ORF1b as a highly derived, inactive N7-MTase-like domain, setting it apart from other N7-MTases in Nidovirales, which however retain catalytic features while sharing the derived Rossmann fold. In non-mammalian arteriviruses, we identified a canonical 2ʹ-O-MTase and an active ExoN domain within ORF1b, both of which are notably absent in their mammalian counterparts. We also noted a probable reduction in PLPs and structural glycoproteins in non-mammalian arteriviruses, emphasizing host-specific divergences in structural proteins. Overall, our results enhance the understanding of the unique features and evolutionary dynamics of mammalian and non-mammalian arteriviruses within nidovirales. We hope that the testable predictions presented here will encourage future experimental investigations of these domains and further our understanding of arterivirus biology.

## Data availability

The data underlying this article are available in the article and in its online supplementary material. This data is also available at https://zenodo.org/records/14172365.

