## Supplementary Material for "Uncovering the Structural and Functional Implications of Uncharacterized NSPs and Variations in the Molecular Toolkit Across Mammalian and Non-Mammalian Arteriviruses"

7. **Supplementary Datasets S2:** Structural proteins phylogenetic tree raw files in Newick format.......**22-25**

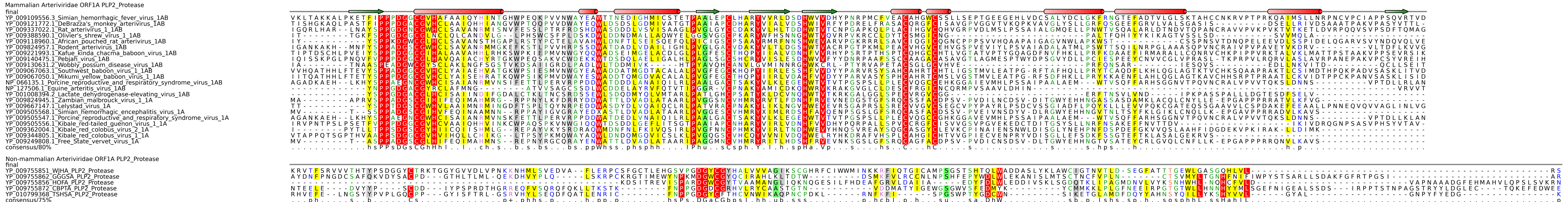
Mammalian Arteriviridae ORF1A PLP2_Protease
final
YP_009109556.3_Simian_hemorrhagic_fever_virus_1AB
YP_009121772.1_DeBrazza's_monkey_arterivirus_1AB
YP_009337021.1_Rat_arterivirus_1_1AB
YP_009388590.1_Olivier's_shrew_virus_1_1AB
YP_009118960.1_African_pouched_rat_arterivirus_1AB
YP_009824957.1_Rodent_arterivirus_1AB
YP_009221993.1_Kafue_kinda_chacma_baboon_virus_1AB
YP_009140475.1_Pebjah_virus_1AB
YP_009130631.2_Wobbly_possum_disease_virus_1AB
YP_009067063.1_Southwest_baboon_virus_1_1AB
YP_009067050.1_Mikumi_yellow_baboon_virus_1_1AB
NP_066135.1_Porcine_reproductive_and_respiratory_syndrome_virus_1AB
NP_127506.1_Equine_arteritis_virus_1AB
YP_001008394.2_Lactate_dehydrogenase-elevating_virus_1AB
YP_009824945.1_Zambian_malbrouck_virus_1_1A
YP_009667147.1_Lelystad_virus_1A
YP_009505568.1_Simian_hemorrhagic_encephalitis_virus_1A
YP_009505547.1_Porcine_reproductive_and_respiratory_syndrome_virus_1A
YP_009505556.1_Kibale_red-tailed_guenon_virus_1_1A
YP_009362004.1_Kibale_red_colobus_virus_2_1A
YP_009344805.1_Kibale_red_colobus_virus_1_1A
YP_009249808.1_Free_State_vervet_virus_1A
consensus/80%
Non-mammalian Arteriviridae ORF1A PLP2_Protease
final
YP_009755851_WJHA_PLP2_Protease
YP_009755862_GGGSA_PLP2_Protease
YP_009755856_HOFA_PLP2_Protease
YP_009755872_CBPTA_PLP2_Protease
YP_010799368_TSHSA_PLP2_Protease
consensus/75%

Mammalian and Non-mammalian Arteriviridae NSP3C wHTH
final

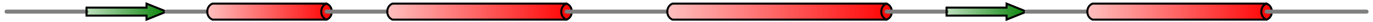

| | |
|---|---|
| YP_009109556.3_Simian_hemorrhagic_fever_virus_1AB | TGIGGFVTPYDIHDVVKSQRDGVAVANAPPNTYLGAVRRAALTGKPAFFVANNTGIVLEGLLREKTRA |
| YP_009388590.1_Olivier's_shrew_virus_1_1AB | RFVPGVITPWDLARFVSNPKNAASVANATPGSYVHALRTSALTGRNMIYIPTTQGITLEGLRRKPLVP |
| YP_009121772.1_DeBrazza's_monkey_arterivirus_1AB | TGTVGLVTPFDIHRLSRSSRDAVAIANAPANTFLGAVRKAALTGKPTYFLADNTGIVLEGLLRHSKPA |
| YP_009118960.1_African_pouched_rat_arterivirus_1AB | VASPALITPHTLGEITNNPLAAMAVANAKPGSYQHTAKTAALTGRCYLHIPSHASVLLEGKLRSKQRS |
| YP_009337022.1_Rat_arterivirus_1_1AB | TQVAGLVTPYDIHSHTSGPRGAAAILTAPEGTYLAAVRRAALTGKTIMFCPSGVGSLLEGAFRTSKPC |
| YP_009824957.1_Rodent_arterivirus_1AB | THVAGLITPYDIHAYTNGPRGASALVTAPEGTYLAAVRRAALTGRTVLYSPSNVGSLLEGAFRTAKPC |
| YP_009221993.1_Kafue_kinda_chacma_baboon_virus_1AB | TGLGGLVTPYDIHLVSKTPRDAIAVANAPPNSYLGAVRMAALTNTNRFYVGSNTGTVLEGLLREKSCA |
| YP_009140475.1_Pebjah_virus_1AB | TGVGGLVTPYDIRNVVKTARDSVAVANAPPNTYLGAVRRAALTGGCVYYIADNTGIVLEGLLREKTPA |
| YP_009130631.2_Wobbly_possum_disease_virus_1AB | SSCGGVVTPTTAAQSAGVPFASLRAAVSRPGTVLKAAYDSAKDGKVRIYIPTGTGVVQEGAMRCNRVP |
| YP_009067063.1_Southwest_baboon_virus_1_1AB | TGVGGLVTPYDIHLCAKTPRDSIAIANAAPNTYLGAVRMAALSGTNKFFIGSNAGIILEGLLREKTKA |
| YP_009067050.1_Mikumi_yellow_baboon_virus_1_1AB | TGVGGLVTPYDIHLCAKTPRDSIAIANAAPNTYLGAVRMAALSGTNRFFVGSNAGIVLEGLLREKTKA |
| NP_066135.1_Porcine_reproductive_and_respiratory_syndrome_virus_1AB | TNVAGLVTPYDIHHYTSGPRGVAALATAPDGTYLAAVRRAALTGRTMLFTPSQLGSLLEGAFRTRKPS |
| NP_127506.1_Equine_arteritis_virus_1AB | SLMGLCVTPAHLARSAGQSREQMVSAAAPTNSLLGVARDCYVTGTTRLYIPKEGGMVFEGLFRSPKAR |
| YP_001008394.2_Lactate_dehydrogenase-elevating_virus_1AB | TSSTGLVTPYDVHLVTSTPRGASSLASAPEGTYLAAVRRSALTGRCCMFVPTNFGSVLEGSLRTRGCA |
| YP_009824945.1_Zambian_malbrouck_virus_1_1A | TGFGGLVTPYDIHAYTTNPRQAIALANAREGTYLASVRQAALTGKPAWFVPSNFGIMVEGALRTGKPH |
| YP_009667147.1_Lelystad_virus_1A | TQVAGIITPYDIHQYTSGPRGAAAVATAPEGTYMAAVRRAALTGRTLIFTPSAVGSLLEGAFRTHKPC |
| YP_009505568.1_Simian_hemorrhagic_encephalitis_virus_1A | TGFGGLVTPYDIHRFSGNPRQAIALANAKEGTYLAAVRQAALTGKPAWFIPSNFGVMVEGALRSGKTH |
| YP_009505547.1_Porcine_reproductive_and_respiratory_syndrome_virus_1A | TNVAGLVTPYDIHHYTSGPRGVAALATAPDGTYLAAVRRAALTGRTMLFTPSQLGSLLEGAFRTRKPS |
| YP_009505556.1_Kibale_red-tailed_guenon_virus_1_1A | TGTLGLVTPYDIHRVVKSPRDSVAIANAPPNTYFGAVRRAALTGKPVLFMADNTGFIFEGAFRNSNCA |
| YP_009362004.1_Kibale_red_colobus_virus_2_1A | TGCCGLVTPYDIHRIVKTSRDAVNVYNAPAGTYLHAVKMAALTNSNKFFIASNAGYVLEGALRSKGRT |
| YP_009344805.1_Kibale_red_colobus_virus_1_1A | TGVGGLVTPYDIHRVVKSSRDAVALANAPHGTYLAAVRDAALSGHNKLFLSNSIGYMLEGALRTGTPA |
| YP_009249808.1_Free_State_vervet_virus_1A | TGFGGLITPYDIHAHTSNPRQAIAVANAKDGTYLAAVRQAALTGKPAWFIPSNFGVMVEGALRTGKPH |
| YP_009755856_wHTH_HOFA | PRCPVYVTPTTLSEISAANGVTLTTAITPEDSLSNCAFRCMMDGKARIFIPKVRNMVQESKARSQTKT |
| AVM87568_wHTH_HHPA | NAAPVFISPSVASSMMSQWGLDLATVAAGPDSVLAALLEASLTGTSVKHTGSPIRYQSRTTLKHHPLS |
| YP_009755872_wHTH_CBPTA | PSAPTVVTPWTSCSLMLKYGVSSSEMCALSSPFIKAALDCSTRGTFSLYKPPTIIPEAAEASRPHNSA |
| consensus/75% | oshsGlVTPaDlp.hspssRsuhulAsAs.soYLuAlRpAALTGps.hahsss.G.hlEGhhRp.p.s |

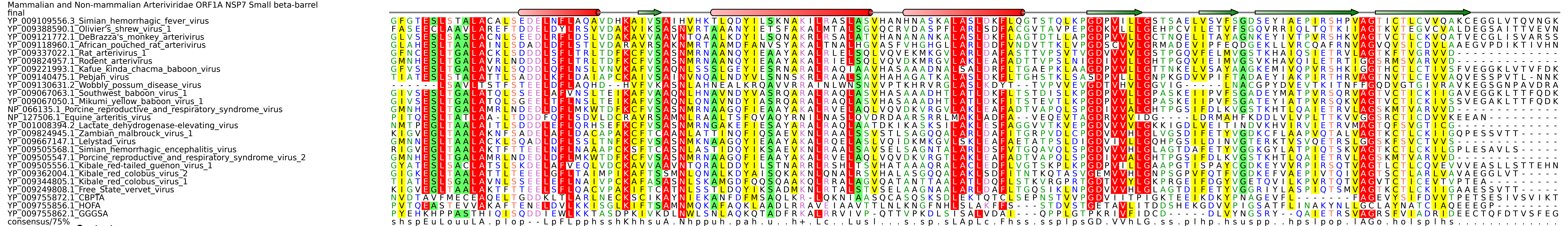
Mammalian and Non-mammalian Arteriviridae ORF1A NSP7 Small beta-barrel
final
YP_009109556.3_Simian_hemorrhagic_fever_virus
YP_009388590.1_Olivier's_shrew_virus_1
YP_009121772.1_DeBrazza's_monkey_arterivirus
YP_009118960.1_African_pouched_rat_arterivirus
YP_009337022.1_Rat_arterivirus_1
YP_009824957.1_Rodent_arterivirus
YP_009221993.1_Kafue_kinda_chacma_baboon_virus
YP_009140475.1_Pebjah_virus
YP_009130631.2_Wobbly_possum_disease_virus
YP_009067063.1_Southwest_baboon_virus_1
YP_009067050.1_Mikumi_yellow_baboon_virus_1
NP_066135.1_Porcine_reproductive_and_respiratory_syndrome_virus
NP_127506.1_Equine_arteritis_virus
YP_001008394.2_Lactate_dehydrogenase-elevating_virus
YP_009824945.1_Zambian_malbrouck_virus_1
YP_009667147.1_Lelystad_virus
YP_009505568.1_Simian_hemorrhagic_encephalitis_virus
YP_009505547.1_Porcine_reproductive_and_respiratory_syndrome_virus_2
YP_009505556.1_Kibale_red-tailed_guenon_virus_1
YP_009362004.1_Kibale_red_colobus_virus_2
YP_009344805.1_Kibale_red_colobus_virus_1
YP_009249808.1_Free_State_vervet_virus
YP_009755872.1_CBPTA
YP_009755856.1_HOFA
YP_009755862.1_GGGSA
consensus/75%

Nidovirales ExoN and Non-mammalian Arterivirus ExoN comparison alignment

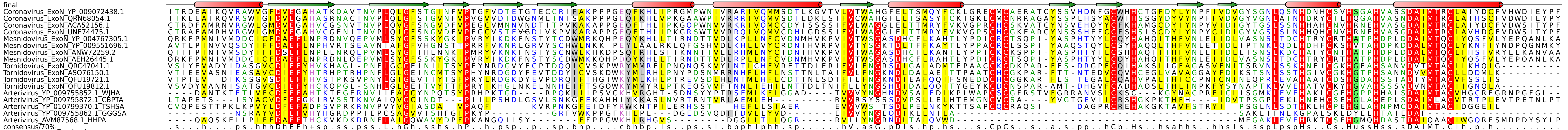

6 Contents

Nidovirales 2'-O-MTase and Non-mammalian Arterivirus 2'-O-MTase comparison alignment

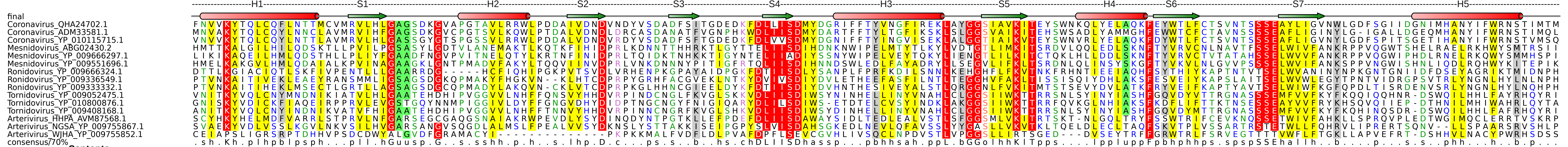

**Contents**

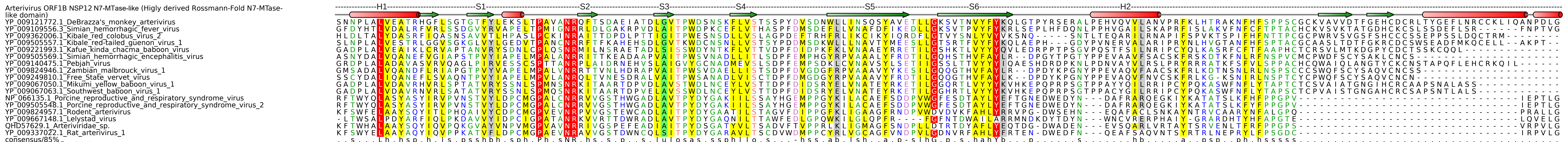
Arterivirus ORF1B NSP12 N7-MTase-like (Higly derived Rossmann-Fold N7-MTase-like domain)
H1
S1
S2
S3
S4
S5
S6
H2
YP_009121772.1_DeBrazza's_monkey_arterivirus
YP_009109556.3_Simian_hemorrhagic_fever_virus
YP_009362006.1_Kibale_red_colobus_virus_2
YP_009505557.1_Kibale_red-tailed_guenon_virus_1
YP_009221993.1_Kafue_kinda_chacma_baboon_virus
YP_009505569.1_Simian_hemorrhagic_encephalitis_virus
YP_009140475.1_Pebjah_virus
YP_009824946.1_Zambian_malbrouck_virus_1
YP_009249810.1_Free_State_vervet_virus
YP_009067050.1_Mikumi_yellow_baboon_virus_1
YP_009067063.1_Southwest_baboon_virus_1
NP_066135.1_Porcine_reproductive_and_respiratory_syndrome_virus
YP_009505548.1_Porcine_reproductive_and_respiratory_syndrome_virus_2
YP_009824957.1_Rodent_arterivirus
YP_009667148.1_Lelystad_virus
QHD57629.1_Arteriviridae_sp.
YP_009337022.1_Rat_arterivirus_1
consensus/85%

Coronavirus ORF1B NSP14-N7-MTase domain (Higly derived Rossmann-Fold; nsp14 : ExoN+N7-MTase)

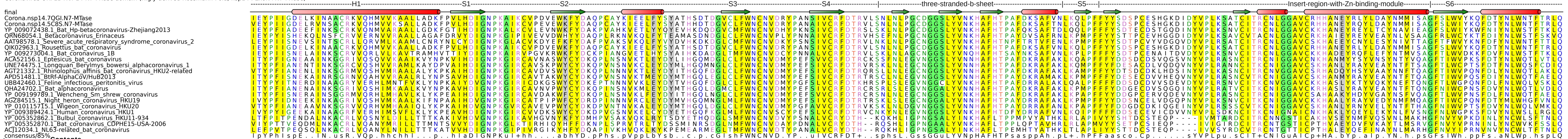

 **Contents**

Mesnidovirus ORF1B N7-MTase alignment (Higly derived Rossmann-Fold)

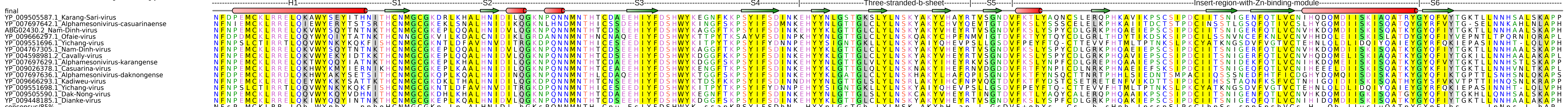

**Contents**

Ronidovirus ORF1B N7-MTase (Higly derived Rossmann-Fold)

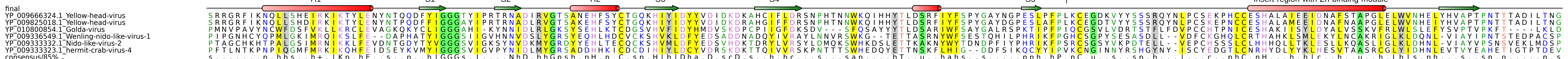

 **Contents**

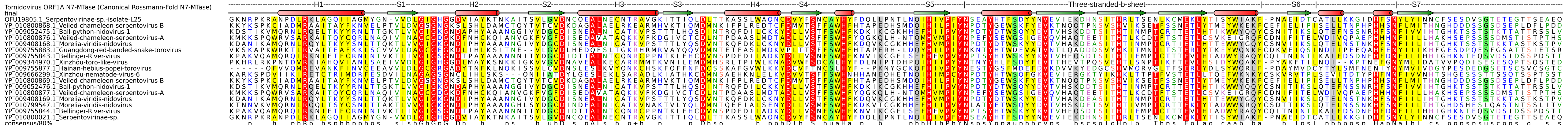

Tornidovirus ORF1A N7-MTase (Canonical Rossmann-Fold N7-MTase)
final
H1
S1
H2
S2
H3
S3
H4
S4
S5
Three-stranded-b-sheet
S6
S7
QFU19805.1_Serpentovirinae-sp.-isolate-L25
YP_010800868.1_Veiled-chameleon-serpentovirus-B
YP_009052475.1_Ball-python-nidovirus-1
YP_010800876.1_Veiled-chameleon-serpentovirus-A
YP_009408168.1_Morelia-viridis-nidovirus
YP_009755883.1_Guangdong-red-banded-snake-torovirus
YP_009755843.1_Bellinger-River-virus
YP_009344970.1_Xinzhou-toro-like-virus
YP_009755877.1_Hainan-hebius-popei-torovirus
YP_009666299.1_Xinzhou-nematode-virus-6
YP_010800869.1_Veiled-chameleon-serpentovirus-B
YP_009052476.1_Ball-python-nidovirus-1
YP_010800877.1_Veiled-chameleon-serpentovirus-A
YP_009408169.1_Morelia-viridis-nidovirus
YP_010799547.1_Morelia-viridis-nidovirus
YP_009755844.1_Bellinger-River-virus
YP_010800021.1_Serpentovirinae-sp.
consensus/80%

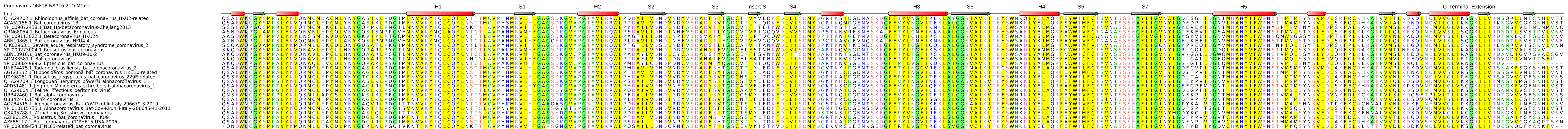
Coronavirus ORF1B NSP16-2'-O-MTase
H1
S1
H2
S2
S3
Insert-S
S4
H3
S5
H4
S6
S7
H5
I
C-Terminal-Extension
final
QHA24702.1_Rhinolophus_affinis_bat_coronavirus_HKU2-related
ACA52156.1_Bat_coronavirus_1B
YP_009072438.1_Bat_Hp-betacoronavirus-Zhejiang2013
QRN68054.1_Betacoronavirus_Erinaceus
YP_009113022.1_Betacoronavirus_HKU24
ABN10865.1_Bat_coronavirus_HKU4-4
QIK02963.1_Severe_acute_respiratory_syndrome_coronavirus_2
YP_009273004.1_Rousettus_bat_coronavirus
ABN10910.1_Bat_coronavirus_HKU9-10-2
ADM33581.1_Bat_coronavirus_HKU9-1
YP_009824989.2_Eptesicus_bat_coronavirus
UNE74475.1_Tadarida_brasiliensis_bat_alphacoronavirus_2
AGT21332.1_Hipposideros_pomona_bat_coronavirus_HKU10-related
UZK98251.1_Rousettus_aegyptiacus_bat_coronavirus_229E-related
QHA24709.1_Longquan_Berylmys_bowersi_alphacoronavirus_1
APD51481.1_Jingmen_Miniopterus_schreibersii_alphacoronavirus_1
QHA24664.1_Feline_infectious_peritonitis_virus
UBB42460.1_Bat_alphacoronavirus
UBB42446.1_Mink_coronavirus_1
AGZ84515.1_Alphacoronavirus_Bat-CoV-P.kuhlii-Italy-206679-3-2010
YP_0101157.1_Alphacoronavirus_Bat-CoV-P.kuhlii-Italy-206645-41-2011
QKX95788.1_Wencheng_Sm_shrew_coronavirus
AZF86129.1_Rousettus_bat_coronavirus_HKU9
AZF86117.1_Bat_coronavirus_CDPHE15-USA-2006
YP_009389424.1_NL63-related_bat_coronavirus
consensus/85%

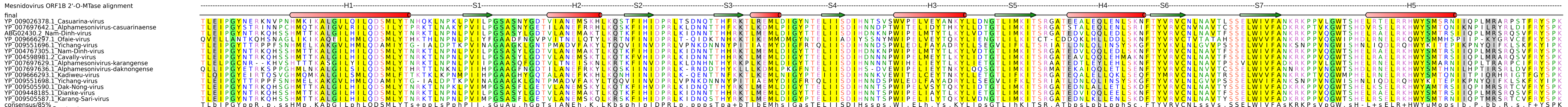
Mesnidovirus ORF1B 2'-O-MTase alignment
final
H1
S1
H2
S2
S3
S4
H3
S5
H4
S6
S7
H5
YP_009026378.1_Casuarina-virus
YP_007697642.1_Alphamesonivirus-casuarinaense
ABG02430.2_Nam-Dinh-virus
YP_009666297.1_Ofaie-virus
YP_009551696.1_Yichang-virus
YP_004767305.1_Nam-Dinh-virus
YP_004598981.2_Cavally-virus
YP_007697629.1_Alphamesonivirus-karangense
YP_007697636.1_Alphamesonivirus-daknongense
YP_009666293.1_Kadiweu-virus
YP_009551698.1_Yichang-virus
YP_009505590.1_Dak-Nong-virus
YP_009448185.1_Dianke-virus
YP_009505587.1_Karang-Sari-virus
consensus/85%
14 **Contents**

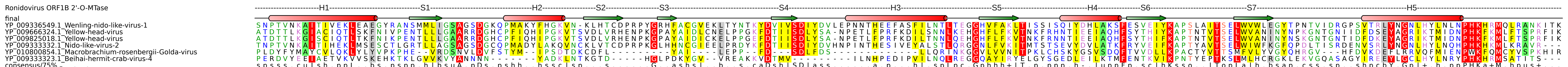

Ronidovirus ORF1B 2'-O-MTase
H1
S1
H2
S2
S3
S4
H3
S5
H4
S6
S7
H5
final
YP_009336549.1_Wenling-nido-like-virus-1
YP_009666324.1_Yellow-head-virus
YP_009825018.1_Yellow-head-virus
YP_009333332.1_Nido-like-virus-2
YP_010800854.1_Macrobrachium-rosenbergii-Golda-virus
YP_009333323.1_Beihai-hermit-crab-virus-4
consensus/75%

Contents

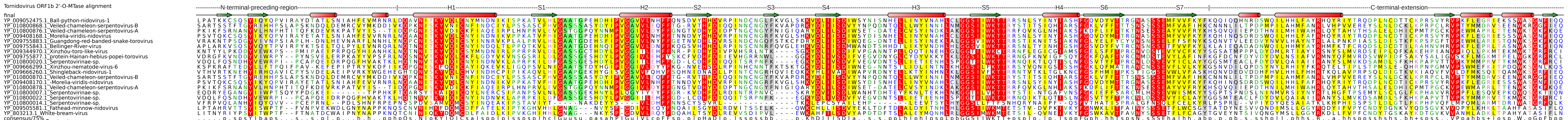

Tornidovirus ORF1b 2'-O-MTase alignment
N-terminal-preceding-region
H1
S1
H2
S2
S3
S4
H3
S5
H4
S6
S7
C-terminal-extension
final
YP_009052475.1_Ball-python-nidovirus-1
YP_010800868.1_Veiled-chameleon-serpentovirus-B
YP_010800876.1_Veiled-chameleon-serpentovirus-A
YP_009408166.1_Morelia-viridis-nidovirus
YP_009755883.1_Guangdong-red-banded-snake-torovirus
YP_009755843.1_Bellinger-River-virus
YP_009344970.1_Xinzhou-toro-like-virus
YP_009755877.1_protein-Hainan-hebius-popei-torovirus
YP_010800020.1_Serpentovirinae-sp.
YP_009666299.1_Xinzhou-nematode-virus-6
YP_009666260.1_Shingleback-nidovirus-1
YP_010800870.1_Veiled-chameleon-serpentovirus-B
YP_009408170.1_Morelia-viridis-nidovirus
YP_010800878.1_Veiled-chameleon-serpentovirus-A
YP_010800071.1_Serpentovirinae-sp.
YP_010800022.1_Serpentovirinae-sp.
YP_010800014.1_Serpentovirinae-sp.
YP_009505581.1_Fathead-minnow-nidovirus
YP_803213.1_White-bream-virus
consensus/75%

 **Contents**

#### Mammalian arterivirus genomes

| # | Genome accession id | Source Organism | Total gene length | CDS_1 | Loci | | CDS_2 | Loci | |
|---|---|---|---|---|---|---|---|---|---|
| 1 | NC_048209.1 | Malbrouck (primate) | 15307 bp | YP_009824945.1<br>2075 aa<br>1A (PLP1α+PLP1β+PLP1γ+PLP2+TM+TM+wHTH+3CL-Pro+TM+Small-β-barrel) | 248-6475 | | YP_009824946.1<br>1467 aa<br>1b (NiRAN+RDRP+ZBD+SF1-Helicase+NendoU+N7-like-MTase) | 6469-10872 | |
| 2 | NC_043487.1 | Porcine (pig) | 15111 bp ss-RNA | YP_009667147.1<br>2396 aa<br>1A (PLP1α+PLP1β+PLP2+TM+TM+wHTH+3CL-Pro+TM+Small-β-barrel) | 222-7412 | | YP_009667148.1<br>1463 aa<br>1b (NiRAN+RDRP+ZBD+SF1-Helicase+NendoU+N7-like-MTase) | 7394-11785 | |
| 3 | NC_038293.1 | simian (primate) | 15370 bp | YP_009505568.1<br>2091 aa<br>1A (PLP1α+PLP1β+PLP1γ+PLP2+TM+TM+wHTH+3CL-Pro+TM+Small-β-barrel) | 205-6480 | | YP_009505569.1<br>1466 aa<br>1b (NiRAN+RDRP+ZBD+SF1-Helicase+NendoU+N7-like-MTase) | 6474-10874 | |
| 4 | NC_038291.1 | Porcine (pig) | 15411 bp | YP_009505547.1<br>2503 aa<br>1A (PLP1α+PLP1β+PLP2+TM+TM+wHTH+3CL-Pro+TM+Small-β-barrel) | 190-7701 | | YP_009505548.1<br>1457 aa<br>1b (NiRAN+RDRP+ZBD+SF1-Helicase+NendoU+N7-like-MTase) | 7698-12071 | 12071-12073 |
| 5 | NC_038292.1 | Guenon (primate) | 15264 bp | YP_009505556.1<br>2060 aa<br>1A (PLP1α+PLP1β+PLP1γ+PLP2+TM+TM+wHTH+3CL-Pro+TM+Small-β-barrel) | 163-6345 | | YP_009505557.1<br>1491 aa<br>1b (NiRAN+RDRP+ZBD+SF1-Helicase+NendoU+N7-like-MTase) | 6339-10814 | |
| 6 | NC_035127.1 | shrew (rodent) | 13766 bp | YP_009388590.1<br>3433 aa<br>1AB (PLP1α+PLP1β+PLP2+TM+TM+wHTH+3CL-Pro+TM+Small-β-barrel+NiRAN+RDRP+ZBD+SF1-Helicase+NendoU+N7-like-MTase) | join(158-6226,6226-10458) | | YP_009388591.1<br>2039 aa<br>1A (PLP1α+PLP1β+PLP2+TM+TM+wHTH+3CL-Pro+TM+Small-β-barrel) | 158-6277 | |
| 7 | NC_026509.1 | Monkey (primate) | 15684 bp | YP_009121772.1<br>3587 aa<br>1AB (PLP1α+PLP1β+PLP1γ+PLP2+TM+TM+wHTH+3CL-Pro+TM+Small-β-barrel+NiRAN+RDRP+ZBD+SF1-Helicase+NendoU+N7-like-MTase) | join(198-6458,6458-10960) | | YP_009121773.1<br>2084 aa<br>1A (PLP1α+PLP1β+PLP1γ+PLP2+TM+TM+wHTH+3CL-Pro+TM+Small-β-barrel) | 210-6464 | |
| 8 | NC_026439.1 | pouched Rat (Rodent) | 14953 bp ss-RNA | YP_009118960.1<br>3714 aa<br>1AB (PLP1α+PLP1β+PLP2+TM+TM+wHTH+3CL-Pro+TM+Small-β-barrel+NiRAN+RDRP+ZBD+SF1-Helicase+NendoU+N7-like-MTase) | join(215-6985,6985-11358) | | YP_009118961.1<br>2240 aa<br>1A (PLP1α+PLP1β+PLP2+TM+TM+wHTH+3CL-Pro+TM+Small-β-barrel) | 269-6991 | |
| 9 | NC_034455.1 | colobus (primate) | 15596 bp | YP_009362004.1<br>2052 aa<br>1A (PLP1α+PLP1β+PLP1γ+PLP2+TM+TM+wHTH+3CL-Pro+TM+Small-β-barrel) | 252-6410 | | YP_009362005.1<br>220 aa<br>nsp2TF | 2776-3438 | |
| 10 | NC_033553.1 | colobus (primate) | 15446 bp | YP_009344805.1<br>2048 aa<br>1A (PLP1α+PLP1β+PLP1γ+PLP2+TM+TM+wHTH+3CL-Pro+TM+Small-β-barrel) | 161-6307 | | YP_009344806.1<br>219 aa<br>nsp2TF | 2724-3383 | |
| 11 | NC_032987.1 | Rat (Rodent) | 15099 bp | YP_009337022.1<br>3925 aa<br>1AB (PLP1α+PLP1β+PLP2+TM+TM+wHTH+3CL-Pro+TM+Small-β-barrel+NiRAN+RDRP+ZBD+SF1-Helicase+NendoU+N7-like-MTase) | | | YP_009337023.1<br>2472 aa<br>1A (PLP1α+PLP1β+PLP2+TM+TM+wHTH+3CL-Pro+TM+Small-β-barrel) | 3-7421 | |
| 12 | NC_029992.1 | Chlorocebus sp. (primate) | 15247 bp | YP_009249808.1<br>2075 aa<br>1A (PLP1α+PLP1β+PLP1γ+PLP2+TM+TM+wHTH+3CL-Pro+TM+Small-β-barrel) | 201-6428 | | YP_009249809.1<br>219 aa<br>nsp2TF | 2836-3495 | |
| 13 | NC_048210.1 | Rodent | 15275 bp | YP_009824957.1<br>3982 aa<br>1AB (PLP1α+PLP1β+PLP2+TM+TM+wHTH+3CL-Pro+TM+Small-β-barrel+NiRAN+RDRP+ZBD+SF1-Helicase+NendoU+N7-like-MTase) | join(<1-7572,7572-11948) | | YP_009824958.1<br>250 aa<br>Glycoprotein 2 | 11924-12676 | |
| 14 | NC_029053.1 | Chicma baboon (primate) | 14924 bp | YP_009221993.1<br>3535 aa<br>1AB (PLP1α+PLP1β+PLP1γ+PLP2+TM+TM+wHTH+3CL-Pro+TM+Small-β-barrel+NiRAN+RDRP+ZBD+SF1-Helicase+NendoU+N7-like-MTase) | join(154-6315,6315-10760) | | YP_009221994.1<br>2055 aa<br>1A (PLP1α+PLP1β+PLP1γ+PLP2+TM+TM+wHTH+3CL-Pro+TM+Small-β-barrel) | 154-6321 | |
| 15 | NC_027124.1 | Macaques (primate) | 15478 bp | YP_009140475.1<br>3552 aa<br>1AB (PLP1α+PLP1β+PLP1γ+PLP2+TM+TM+wHTH+3CL-Pro+TM+Small-β-barrel+NiRAN+RDRP+ZBD+SF1-Helicase+NendoU+N7-like-MTase) | join(165-6347,6347-10822) | | YP_009140476.1<br>2062 aa<br>1A (PLP1α+PLP1β+PLP1γ+PLP2+TM+TM+wHTH+3CL-Pro+TM+Small-β-barrel) | 165-6353 | |
| 16 | NC_026811.2 | Possum (Marsupial) | 12917 bp ss-RNA | YP_009130631.2<br>3402 aa<br>1AB (PLP1α+PLP1β+PLP2+TM+TM+wHTH+3CL-Pro+TM+Small-β-barrel+NiRAN+RDRP+ZBD+SF1-Helicase+NendoU+N7-like-MTase) | join(246-6218,6218-10453) | | YP_009130632.2<br>1998 aa<br>1A (PLP1α+PLP1β+PLP2+TM+TM+wHTH+3CL-Pro+TM+Small-β-barrel) | 246-6242 | |
| 17 | NC_025113.1 | baboon (primate) | 14851 bp | YP_009067063.1<br>3540 aa<br>1AB (PLP1α+PLP1β+PLP1γ+PLP2+TM+TM+wHTH+3CL-Pro+TM+Small-β-barrel+NiRAN+RDRP+ZBD+SF1-Helicase+NendoU+N7-like-MTase) | join(91-6252,6252-10712) | | YP_009067064.1<br>2055 aa<br>1A (PLP1α+PLP1β+PLP1γ+PLP2+TM+TM+wHTH+3CL-Pro+TM+Small-β-barrel) | 91-6258 | |
| 18 | NC_025112.1 | Baboon (primate) | 14927 bp | YP_009067050.1<br>3541 aa<br>1AB (PLP1α+PLP1β+PLP1γ+PLP2+TM+TM+wHTH+3CL-Pro+TM+Small-β-barrel+NiRAN+RDRP+ZBD+SF1-Helicase+NendoU+N7-like-MTase) | join(166-6330,6330-10790) | | YP_009067051.1<br>2056 aa<br>1A (PLP1α+PLP1β+PLP1γ+PLP2+TM+TM+wHTH+3CL-Pro+TM+Small-β-barrel) | 166-6336 | |
| 19 | NC_001961.1 | Porcine (Pig) | 15428 bp ss-RNA | NP_066135.1<br>3960 aa<br>1AB (PLP1α+PLP1β+PLP2+TM+TM+wHTH+3CL-Pro+TM+Small-β-barrel+NiRAN+RDRP+ZBD+SF1-Helicase+NendoU+N7-like-MTase) | join(190-7695,7695-12071) | | NP_047406.1<br>2503 aa<br>1A (PLP1α+PLP1β+PLP2+TM+TM+wHTH+3CL-Pro+TM+Small-β-barrel) | 190-7701 | |
| 20 | NC_002532.2 | Equine (Horse) | 12704 bp ss-RNA | NP_127506.1<br>3175 aa<br>1AB (PLP1α+PLP1β+PLP2+TM+TM+wHTH+3CL-Pro+TM+Small-β-barrel+NiRAN+RDRP+ZBD+SF1-Helicase+NendoU+N7-like-MTase) | join(225-5405,5405-9751) | | NP_127507.1<br>1727 aa<br>1A (PLP1α+PLP1β+PLP2+TM+TM+wHTH+3CL-Pro+TM+Small-β-barrel) | 225-5408 | |
| 21 | NC_003092.2 | Simian (primate) | 15717 bp | YP_009109556.3<br>3595 aa<br>1AB (PLP1α+PLP1β+PLP1γ+PLP2+TM+TM+wHTH+3CL-Pro+TM+Small-β-barrel+NiRAN+RDRP+ZBD+SF1-Helicase+NendoU+N7-like-MTase) | join(210-6521,6521-10996) | | NP_203542.2<br>2105 aa<br>1A (PLP1α+PLP1β+PLP1γ+PLP2+TM+TM+wHTH+3CL-Pro+TM+Small-β-barrel) | 210-6527 | |
| 22 | NC_001639.1 | Environment | 14104 bp ss-RNA | YP_001008394.2<br>3616 aa<br>1AB (PLP1α+PLP1β+PLP2+TM+TM+wHTH+3CL-Pro+TM+Small-β-barrel+NiRAN+RDRP+ZBD+SF1-Helicase+NendoU+N7-like-MTase) | join(157-6771,6771-11006) | | NP_042572.1<br>2206 aa<br>1A (PLP1α+PLP1β+PLP2+TM+TM+wHTH+3CL-Pro+TM+Small-β-barrel) | 157-6777 | |

| # | CDS_3 | Loci | | CDS_4 | Loci | | CDS_5 | Loci | | CDS_6 | Loci | | CDS_7 | Loci | | CDS_8 | Loci |
|---|---|---|---|---|---|---|---|---|---|---|---|---|---|---|---|---|---|
| 1 | YP_009824947.1<br>237 aa<br>Glycoprotein 2' | 10806-11519 | | YP_009824948.1<br>199 aa<br>Glycoprotein 3' | 11354-11953 | | YP_009824949.1<br>170 aa<br>Glycoprotein 4' | 11878-12390 | | YP_009824950.1<br>81 aa<br>E_protein | 12383-12628 | | YP_009824951.1<br>190 aa<br>Glycoprotein 2 | 12423-12995 | | YP_009824952.1<br>270 aa<br>Glycoprotein 3 | 12884-13696 |
| 2 | YP_009667149.1<br>249 aa<br>Glycoprotein 2 | 11796-12545 | | YP_009667150.1<br>265 aa<br>Glycoprotein 3 | 12404-13201 | | YP_009667151.1<br>183 aa<br>Glycoprotein 4 | 12946-13497 | | YP_009667152.1<br>201 aa<br>Glycoprotein 5 | 13494-14099 | | YP_009667153.1<br>173 aa<br>M | 14087-14608 | | YP_009667154.1<br>128 aa<br>N | 14598-14984 |
| 3 | YP_009505570.1<br>235 aa<br>Glycoprotein 2' | 10811-11518 | | YP_009505571.1<br>199 aa<br>Glycoprotein 3' | 11353-11952 | | YP_009505572.1<br>170 aa<br>Glycoprotein 4' | 11877-12389 | | YP_009505573.1<br>81 aa<br>E_protein | 12382-12627 | | YP_009505574.1<br>190 aa<br>Glycoprotein 2 | 12422-12994 | | YP_009505575.1<br>156 aa<br>Glycoprotein 3 | 12883-13353 |
| 4 | YP_009505549.1<br>256 aa<br>Glycoprotein 2 | 12073-12843 | | YP_009505550.1<br>73 aa<br>E_protein | 12078-12299 | | YP_009505551.1<br>254 aa<br>Glycoprotein 3 | 12696-13460 | | YP_009505552.1<br>178 aa<br>Glycoprotein 4 | 13241-13777 | 13777-13788 | YP_009505553.1<br>200 aa<br>Glycoprotein 5 | 13788-14390 | | YP_009505554.1<br>174 aa<br>M | 14375-14899 |
| 5 | YP_009505558.1<br>214 aa<br>Glycoprotein 2' | 10805-11449 | | YP_009505559.1<br>207 aa<br>Glycoprotein 3' | 11320-11943 | | YP_009505560.1<br>198 aa<br>Glycoprotein 4' | 11805-12401 | | YP_009505561.1<br>79 aa<br>E_protein | 12394-12633 | | YP_009505562.1<br>190 aa<br>Glycoprotein 2 | 12434-13006 | | YP_009505563.1<br>136 aa<br>Glycoprotein 3 | 12856-13266 |
| 6 | YP_009388592.1<br>71 aa<br>E_protein | 10451-10666 | | YP_009388593.1<br>300 aa<br>Glycoprotein 2 | 10479-11381 | | YP_009388594.1<br>183 aa<br>Glycoprotein 3 | 11216-11767 | | YP_009388595.1<br>157 aa<br>Glycoprotein 4 | 11686-12159 | | YP_009388596.1<br>206 aa<br>Glycoprotein 5 | 12156-12776 | | YP_009388597.1<br>48 aa<br>Glycoprotein 5a | 12161-12307 |
| 7 | YP_009121774.1<br>214 aa<br>Glycoprotein 2' | 10939-11583 | | YP_009121775.1<br>213 aa<br>Glycoprotein 3' | 11454-12095 | | YP_009121776.1<br>202 aa<br>Glycoprotein 4' | 11951-12559 | | YP_009121777.1<br>235 aa<br>Glycoprotein 2 | 12505-13212 | | YP_009121778.1<br>80 aa<br>E_protein | 12552-12794 | | YP_009121779.1<br>166 aa<br>Glycoprotein 3 | 13083-13583 |
| 8 | YP_009118962.1<br>74 aa<br>E_protein | 11337-11561 | | YP_009118963.1<br>304 aa<br>Glycoprotein 2 | 11368-12282 | | YP_009118964.1<br>245 aa<br>Glycoprotein 3 | 12033-12770 | | YP_009118965.1<br>180 aa<br>Glycoprotein 4 | 12668-13210 | | YP_009118966.1<br>229 aa<br>Glycoprotein 5 | 13138-13827 | | YP_009118967.1<br>59 aa<br>Glycoprotein 5a | 13200-13379 |
| 9 | YP_009362006.1<br>1485 aa<br>1b (NiRAN+RDRP+ZBD+SF1-Helicase+NendoU+N7-like-MTase) | 6404-10861 | | YP_009362007.1<br>223 aa<br>Glycoprotein 2' | 10744-11415 | | YP_009362008.1<br>202 aa<br>Glycoprotein 3' | 11286-11894 | | YP_009362009.1<br>166 aa<br>Glycoprotein 4' | 11819-12319 | | YP_009362010.1<br>81 aa<br>E_protein | 12312-12557 | | YP_009362011.1<br>204 aa<br>Glycoprotein 2 | 12352-12966 |
| 10 | YP_009344807.1<br>1464 aa<br>1b (NiRAN+RDRP+ZBD+SF1-Helicase+NendoU+N7-like-MTase) | 6301-10695 | | YP_009344808.1<br>232 aa<br>Glycoprotein 2' | 10635-11333 | | YP_009344809.1<br>220 aa<br>Glycoprotein 3' | 11186-11848 | | YP_009344810.1<br>194 aa<br>Glycoprotein 4' | 11773-12357 | | YP_009344811.1<br>80 aa<br>E_protein | 12345-12587 | | YP_009344812.1<br>205 aa<br>Glycoprotein 2 | 12385-13002 |
| 11 | YP_009337024.1<br>70 aa<br>E_protein | 11789-12001 | | YP_009337025.1<br>254 aa<br>Glycoprotein 2 | 11829-12593 | 12597-12607 | YP_009337026.1<br>191 aa<br>Glycoprotein 3 | 12607-13182 | | YP_009337027.1<br>182 aa<br>Glycoprotein 4 | 12933-13481 | | YP_009337028.1<br>45 aa<br>Glycoprotein 5a | 13483-13620 | | YP_009337029.1<br>193 aa<br>Glycoprotein 5 | 13511-14092 |
| 12 | YP_009249810.1<br>1465 aa<br>1b (NiRAN+RDRP+ZBD+SF1-Helicase+NendoU+N7-like-MTase) | 6422-10819 | | YP_009249811.1<br>236 aa<br>Glycoprotein 2' | 10756-11466 | | YP_009249812.1<br>199 aa<br>Glycoprotein 3' | 11301-11900 | | YP_009249813.1<br>170 aa<br>Glycoprotein 4' | 11825-12337 | | YP_009249814.1<br>81 aa<br>E_protein | 12330-12575 | | YP_009249815.1<br>190 aa<br>Glycoprotein 2 | 12370-12942 |
| 13 | YP_009824959.1<br>70 aa<br>E_protein | 11932-12144 | | YP_009824960.1<br>265 aa<br>Glycoprotein 3 | 12535-13332 | | YP_009824961.1<br>182 aa<br>Glycoprotein 4 | 13077-13625 | | YP_009824962.1<br>203 aa<br>Glycoprotein 5 | 13622-14233 | | YP_009824963.1<br>174 aa<br>M | 14218-14742 | | YP_009824964.1<br>N | 14732-15103 |
| 14 | YP_009221995.1<br>1060 aa<br>TF | join(154-2681,2680-3334) | | YP_009221996.1<br>220 aa<br>Glycoprotein 2' | 10714-11376 | | YP_009221997.1<br>202 aa<br>Glycoprotein 3' | 11250-11858 | | YP_009221998.1<br>157 aa<br>Glycoprotein 4' | 11783-12256 | | YP_009221999.1<br>75 aa<br>E_protein | 12249-12476 | | YP_009222000.1<br>158 aa<br>Glycoprotein 2 | 12277..12753 |
| 15 | YP_009140477.1<br>3205<br>TF | join(165-2687,2689-3366) | | YP_009140478.1<br>223 aa<br>Glycoprotein 2' | 10746-11417 | | YP_009140479.1<br>52 aa<br>2b' | 10852-11010 | | YP_009140480.1<br>213 aa<br>Glycoprotein 3' | 11288-11929 | | YP_009140481.1<br>175 aa<br>Glycoprotein 4' | 11854-12381 | | YP_009140482.1<br>80 aa<br>E_protein | 12374-12616 |
| 16 | YP_009130633.1<br>207 aa<br>Glycoprotein 2 | 10309-10932 | | YP_009130634.1<br>45 aa<br>E_protein | 10553-10690 | | YP_009130635.1<br>217 aa<br>Glycoprotein 3 | 10794-11447 | | YP_009130636.1<br>83 aa<br>Glycoprotein 4 | 11330-11581 | | YP_009130637.1<br>178 aa<br>Glycoprotein 5 | 11578-12114 | | YP_009130638.1<br>78 aa<br>Glycoprotein 5a | 11603-11839 |
| 17 | YP_009067065.1<br>222 aa<br>Glycoprotein 2' | 10660-11328 | | YP_009067066.1<br>52 aa<br>2b' | 10766-10924 | | YP_009067067.1<br>201 aa<br>Glycoprotein 3' | 11202-11807 | | YP_009067068.1<br>157 aa<br>Glycoprotein 4' | 11732-12205 | | YP_009067069.1<br>76 aa<br>E_protein | 12198-12428 | | YP_009067070.1<br>173 aa<br>Glycoprotein 2 | 12208-12729 |
| 18 | YP_009067052.1<br>222 aa<br>Glycoprotein 2' | 10738-11406 | | YP_009067053.1<br>202 aa<br>Glycoprotein 3' | 11280-11888 | | YP_009067054.1<br>157 aa<br>Glycoprotein 4' | 11813-12286 | | YP_009067055.1<br>76 aa<br>E_protein | 12279-12509 | | YP_009067056.1<br>175 aa<br>Glycoprotein 2 | 12289-12816 | | YP_009067057.1<br>144 aa<br>Glycoprotein 3 | 12597-13031 |
| 19 | YP_006950436.1<br>1402 aa<br>TF | join(190-3888,3887-4396) | | NP_047408.1<br>256 aa<br>Glycoprotein 2 | 12073-12843 | | YP_006589918.1<br>73 aa<br>E_protein | 12078-12299 | | NP_047409.1<br>254 aa<br>Glycoprotein 3 | 12696-13460 | | NP_047410.1<br>178 aa<br>Glycoprotein 4 | 13241-13777 | | YP_006488613.1<br>51 aa<br>Glycoprotein 5a | 13778-13933 |
| 20 | NP_127508.1<br>67 aa<br>E_protein | 9751-9954 | | NP_065656.1<br>227 aa<br>Glycoprotein 2 | 9824-10507 | | NP_065657.1<br>163 aa<br>Glycoprotein 3 | 10306-10797 | | NP_065658.1<br>152 aa<br>Glycoprotein 4 | 10700-11158 | | YP_006390077.1<br>59 aa<br>Glycoprotein 5a | 11112-11291 | | NP_065659.1<br>255 aa<br>Glycoprotein 5 | 11146-11913 |
| 21 | YP_009172490.1<br>1114 aa<br>TF | join(210-2876,2875-3552) | | NP_203544.1<br>281 aa<br>Glycoprotein 2' | 10953-11798 | | YP_009037599.1<br>94 aa<br>2b' | 11047-11331 | | NP_203545.1<br>204 aa<br>Glycoprotein 3' | 11486-12100 | | NP_203546.1<br>205 aa<br>Glycoprotein 4' | 11938-12555 | | YP_009037600.1<br>80 aa<br>E_protein | 12555-12797 |
| 22 | YP_006950434.1<br>1117 aa<br>TF | join(157-3000,2999-3508) | | YP_006589917.1<br>70 aa<br>E_protein | 11006-11218 | | NP_042574.1<br>227 aa<br>Glycoprotein 2 | 11046-11729 | | NP_042575.1<br>191 aa<br>Glycoprotein 3 | 11597-12172 | | NP_042576.1<br>175 aa<br>Glycoprotein 4 | 12064-12591 | | NP_042577.1<br>199 aa<br>Glycoprotein 5 | 12588-13187 |

| # | CDS_9 | Loci | | CDS_10 | Loci | | CDS_11 | Loci | | CDS_12 | Loci | | CDS_13 | Loci | | CDS_14 | Loci | | CDS_15 | Loci |
|---|---|---|---|---|---|---|---|---|---|---|---|---|---|---|---|---|---|---|---|---|
| 1 | YP_009824953.1<br>59 aa<br>Glycoprotein 5a | 13689-13868 | | YP_009824954.1<br>241 aa<br>Glycoprotein 5 | 13732-14457 | | YP_009824955.1<br>159 aa<br>M | 14451-14930 | | YP_009824956.1<br>102 aa<br>N | 14911-15219 | | | | | | | | | |
| 2 | | | | | | | | | | | | | | | | | | | | |
| 3 | YP_009505576.1<br>174 aa<br>Glycoprotein 4 | 13176-13700 | | YP_009505577.1<br>59 aa<br>Glycoprotein 5a | 13693-13872 | | YP_009505578.1<br>Glycoprotein 5 | 13736-14479 | | YP_009505579.1<br>159 aa<br>M/GP | 14473-14952 | | YP_009505580.1<br>108 aa<br>N | 14933-15259 | | | | | | |
| 4 | YP_009505555.1<br>123 aa<br>N | 14889-15260 | | | | | | | | | | | | | | | | | | |
| 5 | YP_009505564.1<br>175 aa<br>Glycoprotein 4 | 13074-13601 | 13601-13604 | YP_009505565.1<br>261 aa<br>Glycoprotein 5 | | | YP_009505566.1<br>161 aa<br>M | 14383-14868 | | YP_009505567.1<br>131 aa<br>N | 14798-15193 | | | | | | | | | |
| 6 | YP_009388598.1<br>171 aa<br>M | 12779-13294 | | YP_009388599.1<br>121 aa<br>N | 13287-13652 | | | | | | | | | | | | | | | |
| 7 | YP_009121780.1<br>179 aa<br>Glycoprotein 4 | 13442-13981 | | YP_009121781.1<br>265 aa<br>Glycoprotein 5 | 13984-14781 | | YP_009121782.1<br>59 aa<br>Glycoprotein 5a | 13989-14168 | | YP_009121783.1<br>173 aa<br>M | 14739-15260 | | YP_009121784.1<br>115 aa<br>N | 15235-15582 | | | | | | |
| 8 | YP_009118968.1<br>173 aa<br>M | 13815-14336 | | YP_009118969.1<br>134 aa<br>N | 14333-14737 | | | | | | | | | | | | | | | |
| 9 | YP_009362012.1<br>195 aa<br>Glycoprotein 3 | 12846-13433 | | YP_009362013.1<br>171 aa<br>Glycoprotein 4 | 13337-13852 | | YP_009362014.1<br>58 aa<br>Glycoprotein 5a | 13852-14028 | | YP_009362015.1<br>232 aa<br>Glycoprotein 5 | 13904-14602 | | YP_009362016.1<br>160 aa<br>M | 14596-15078 | | YP_009362017.1<br>111 aa<br>N | 15056-15391 | | | |
| 10 | YP_009344813.1<br>199 aa<br>Glycoprotein 3 | 12801-13400 | | YP_009344814.1<br>173 aa<br>Glycoprotein 4 | 13247-13768 | | YP_009344815.1<br>68 aa<br>Glycoprotein 5a | 13768-13974 | | YP_009344816.1<br>246 aa<br>Glycoprotein 5 | 13820-14560 | | YP_009344817.1<br>161 aa<br>M | 14554-15039 | | YP_009344818.1<br>119 aa<br>N | 15014-15373 | | | |
| 11 | YP_009337030.1<br>174 aa<br>M | 14077-14601 | | YP_009337031.1<br>122 aa<br>N | 14591-14959 | | | | | | | | | | | | | | | |
| 12 | YP_009249816.1<br>155 aa<br>Glycoprotein 3 | 12831-13298 | | YP_009249817.1<br>170 aa<br>Glycoprotein 4 | 13133-13645 | | YP_009249818.1<br>59 aa<br>Glycoprotein 5a | 13635-13814 | | YP_009249819.1<br>244 aa<br>Glycoprotein 5 | 13678-14412 | | YP_009249820.1<br>159 aa<br>M | 14406-14885 | | YP_009249821.1<br>111 aa<br>N | 14866-15201 | | | |
| 13 | | | | | | | | | | | | | | | | | | | | |
| 14 | YP_009222001.1<br>146 aa<br>Glycoprotein 3 | 12558..12998 | | YP_009222002.1<br>152 aa<br>Glycoprotein 4 | 12827-13285 | | YP_009222003.1<br>268 aa<br>Glycoprotein 5 | 13288-14094 | | YP_009222004.1<br>50 aa<br>Glycoprotein 5a | 13296-13448 | | YP_009222005.1<br>161 aa<br>M | 14088-14573 | | YP_009222006.1<br>111 aa<br>N | 14551-14886 | | | |
| 15 | YP_009140483.1<br>202 aa<br>Glycoprotein 2 | 12414-13022 | | YP_009140484.1<br>178 aa<br>Glycoprotein 3 | 12893-13429 | | YP_009140485.1<br>176 aa<br>Glycoprotein 4 | 13270-13800 | 13800-13814 | YP_009140486.1<br>61 aa<br>Glycoprotein 5a | 13814-13999 | | YP_009140487.1<br>248 aa<br>Glycoprotein 5 | 13866-14612 | | YP_009140488.1<br>161 aa<br>M | 14606-15091 | | YP_009140489.1<br>115 aa<br>N | 15066-15413 |
| 16 | YP_009130639.1<br>180 aa<br>M | 12051-12593 | | YP_009130640.1<br>126 aa<br>N | 12424-12804 | | | | | | | | | | | | | | | |
| 17 | YP_009067071.1<br>143 aa<br>Glycoprotein 3 | 12516-12947 | | YP_009067072.1<br>158 aa<br>Glycoprotein 4 | 12767-13243 | | YP_009067073.1<br>258 aa<br>Glycoprotein 5 | 13236-14012 | | YP_009067074.1<br>48 aa<br>Glycoprotein 5a | 13244-13390 | | YP_009067075.1<br>161 aa<br>M | 14006-14491 | | YP_009067076.1<br>115 aa<br>N | 14469-14816 | | | |
| 18 | YP_009067058.1<br>158 aa<br>Glycoprotein 4 | 12848-13324 | | YP_009067059.1<br>256 aa<br>Glycoprotein 5 | 13317-14087 | | YP_009067060.1<br>49 aa<br>Glycoprotein 5a | 13325-13474 | | YP_009067061.1<br>161 aa<br>M | 14081-14566 | | YP_009067062.1<br>115 aa<br>N | 14544-14891 | | | | | | |
| 19 | NP_047411.1<br>200 aa<br>Glycoprotein 5 | 13788-14390 | | NP_047412.1<br>174 aa<br>M | 14375-14899 | | NP_047413.1<br>123 aa<br>N | 14889-15260 | | | | | | | | | | | | |
| 20 | NP_065660.1<br>162 aa<br>M | 11901-12389 | | NP_065661.1<br>110 aa<br>N | 12313-12645 | | | | | | | | | | | | | | | |
| 21 | NP_203547.1<br>214 aa<br>Glycoprotein 2 | 12595-13239 | | NP_203548.1<br>179 aa<br>Glycoprotein 3 | 13089-13628 | | NP_203549.1<br>182 aa<br>Glycoprotein 4 | 13451-13999 | 13999-14002 | NP_203550.2<br>278 aa<br>Glycoprotein 5 | 14002-14838 | | YP_009037601.1<br>64 aa<br>Glycoprotein 5a | 14010-14204 | | NP_203551.1<br>162 aa<br>M | 14832-15320 | | NP_203552.1<br>111 aa<br>N | 15295-15630 |
| 22 | YP_006488612.1<br>47 aa<br>Glycoprotein 5a | 12593-12736 | | NP_042578.1<br>171 aa<br>M | 13175-13690 | | NP_042579.1<br>115 aa<br>N | 13677-14024 | | | | | | | | | | | | |

#### Non-mammalian arterivirus genomes

| # | Genome accession id | Source Organism | Total gene length | CDS_1 | Loci | | CDS_2 | Loci |
|---|---|---|---|---|---|---|---|---|
| 1 | NC_046957.1 | Hyporhamphus sajori<br>Actinopterygii<br>Wuhan japanese halfbeak arterivirus (WJHA) | 18239 bp | YP_009755851.1<br>2196 aa<br>ORF1A<br>DIV-1 + PLP2_Protease + DIV-2 + 4TM + Alpha+Beta + DIV-3 + 5TM + 3CL_Protease + 5TM + Alpha+Beta | 993-7583 | 7583-7880 | YP_009755852.1<br>2864 aa<br>ORF1B<br>DIV-4 + NiRAN+RDRP + ZFD+SF1-Helicase + ExoN(DEDDh) + NendoU + 2'-O-MTase + DIV-5 | 7880-16474 |
| 2 | NC_046960.1 | Chimaera sp<br>Chondrichthyes<br>Nanhai ghost shark arterivirus (NGSA) | 13162 bp | YP_009755867.1<br>3399 aa<br>ORF1AB<br>DIV-1 + 5TM + Beta_rich + 4TM + 3CL_Protease + 2TM + Alpha+Beta + NiRAN+RDRP + ZFD+SF1-Helicase + NendoU + 2'-O-MTase + DIV-2 | join(670-5169,5169-10868) | | YP_009755868.1<br>305 aa<br>GP-like fold | 10216-11133 |
| 3 | NC_046959.1 | Cyclophiops major<br>Reptilia-snake<br>Guangdong greater green snake arterivirus (GGGSA) | 18410 bp | YP_009755862.1<br>4326 aa<br>ORF1AB<br>Alpha+Beta + DIV-1 + PLP2_Protease + DIV-2 + 3TM + Alpha_Helical + 7TM + 3CL_Protease + 4TM + Small_Beta-barrel + NiRAN+RDRP + ZFD+SF1-Helicase + ExoN(DEDDh) + NendoU | join(1180-9093,9093-14159) | | YP_009755863.1<br>799 aa<br>GP-like fold | 14205-16604 |
| 4 | NC_046958.1 | Oligodon formosanus<br>Reptilia-snake<br>Hainan oligodon formosanus arterivirus (HOFA) | 15301 bp | YP_009755856.1<br>3821 aa<br>ORF1AB<br>PLP1_Protease + DIV-1 + PLP2_Protease + 4TM + DIV-2 + 6TM + wHTH + 3CL_Protease + 2TM + Small_Beta-barrel + NiRAN+RDRP + ZFD+SF1-Helicase + Alpha+Beta + NendoU + Alpha+Beta | 76-12240 | | YP_009755857.1<br>334 aa<br>GP-like fold | 12231-13235 |
| 5 | MG600021.1 | Hebius popei<br>Reptilia-snake<br>Hainan hebius popei arterivirus (HHPA) | 18239 bp | AVM87568.1<br>3083 aa<br>ORF1AB<br>Alpha+Beta + 4TM + wHTH + 3CL_Protease + 6TM + Alpha+Beta + NiRAN+RDRP + ZFD+SF1-Helicase + ExoN(DEDDh) + NendoU + 2'-O-MTase | join(<1..3332,3332..9253) | | AVM87569.1<br>568 aa<br>GP-like fold | 9144..10850 |
| 6 | NC_046961.1 | Mauremys megalocephala<br>Chinese broad-headed pond turtle arterivirus (CBPTA) | 14939 bp | YP_009755872.1<br>3511 aa<br>ORF1AB<br>Alpha+Beta + PLP2_Protease + DIV-1 + 4TM + DIV-2 + 4TM + wHTH + 3CL_Protease + 4TM + Small_Beta-barrel + NiRAN+RDRP + ZFD+SF1-Helicase + ExoN(DEDDh) + NendoU + Alpha+Beta | join(461-5929,5929-10995) | | YP_009755873.1<br>119 aa<br>GP? | 10992-11351 |
| 7 | NC_076637.1 | Trionyx sinensis (chinese softshell turtle)<br>Trionyx sinensis hemorrhagic syndrome arterivirus (TSHSA) | 17738 bp | YP_010799367.1<br>78 aa<br>1A (Transmembrane-Segments) | 1187..1423 | | YP_010799368.1<br>307 aa<br>ORF2 (ORF1A equivalent) PLP2_protease | 1423..2346 |

| # | CDS_3 | Loci | | CDS_4 | Loci | | CDS_5 | Loci | | CDS_6 | Loci | | CDS_7 | Loci | | CDS_8 | Loci |
|---|---|---|---|---|---|---|---|---|---|---|---|---|---|---|---|---|---|
| 1 | YP_009755853.1<br>35 aa<br>GP? | 16534-16641 | 16641-16847 | YP_009755854.1<br>214 aa<br>M-protein | 16847-17491 | | YP_009755855.1<br>125 aa<br>Nucleoprotein | 17494-17871 | 17871:18239 bp | | | | | | | | |
| 2 | YP_009755869.1<br>233 aa<br>M-protein | 11660-12361 | | YP_009755870.1<br>107 aa<br>Nucleoprotein | 12361-12684 | | YP_009755871.1<br>207 aa<br>GP? | 12452-13075 | 13075:13162 bp | | | | | | | | |
| 3 | YP_009755864.1<br>327 aa<br>GP-like fold | 16451-17434 | | YP_009755865.1<br>150 aa<br>GP? | 17365-17817 | | YP_009755866.1<br>160 aa<br>Nucleoprotein | 17541-18023 | 18023:18410 bp | | | | | | | | |
| 4 | YP_009755858.1<br>304 aa<br>GP-like fold | 12997-13911 | | YP_009755859.1<br>194 aa<br>M-protein | 13848-14432 | | YP_009755860.1<br>164 aa<br>GP? | 14332-14826 | 14826:15301 bp | YP_009755861.1<br>90 aa<br>GP? | 14777-15049 | | | | | | |
| 5 | AVM87570.1<br>287 aa<br>M-protein | 10874..11737 | | AVM87571.1<br>96 aa<br>GP? | 11442..11732 | | AVM87572.1<br>126 aa<br>Nucleoprotein | 11739..12119 | | | | | | | | | |
| 6 | YP_009755874.1<br>116 aa<br>Nucleoprotein | 11156-11506 | | YP_009755875.1<br>224 aa<br>M-protein | 11515-12189 | 12189-12366 | YP_009755876.1<br>766 aa<br>GP-like fold | 12366-14666 | 14666:14939 bp | | | | | | | | |
| 7 | YP_010799369.1<br>1551 aa<br>ORF1A<br>DIV-1 + 5TM + 3CL_Protease + 4TM + Alpha+Beta | 2309..6964 | | YP_010799370.1<br>1,691 aa<br>ORF1B<br>NiRAN+RDRP + ZFD+SF1-Helicase + ExoN (DEDDh) + NendoU + DIV-2 | 7057..12132 | | YP_010799371.1<br>168 aa<br>GP? | 12129..12635 | | YP_010799372.1<br>193 aa<br>M-protein | 12637..13218 | | YP_010799373.1<br>328 aa<br>GP-like fold | 13391..14377 | | YP_010799374.1<br>713 aa<br>GP-like fold | 14769..16910 |

### Supplementary Table S1

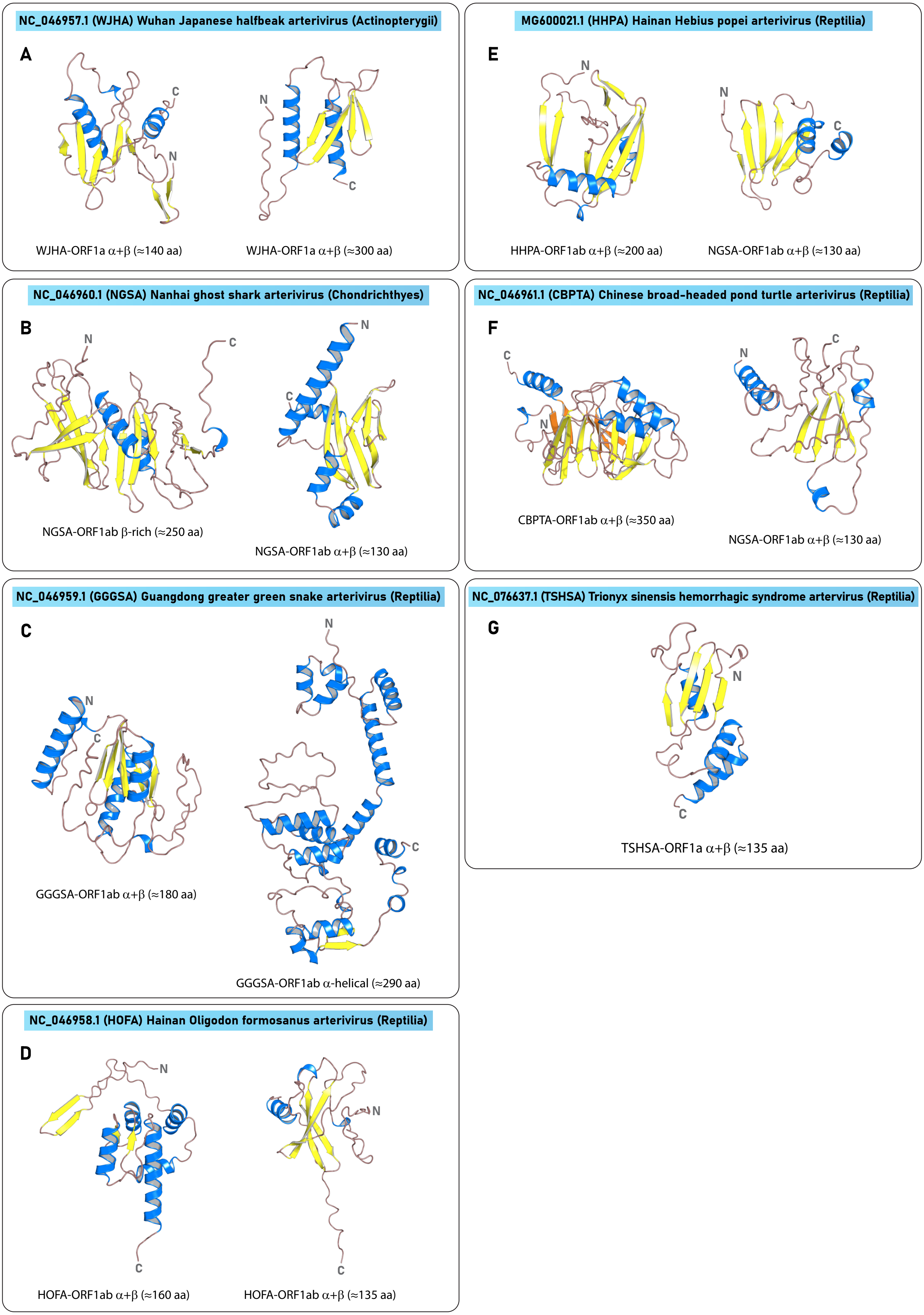

### Supplementary Figure S1

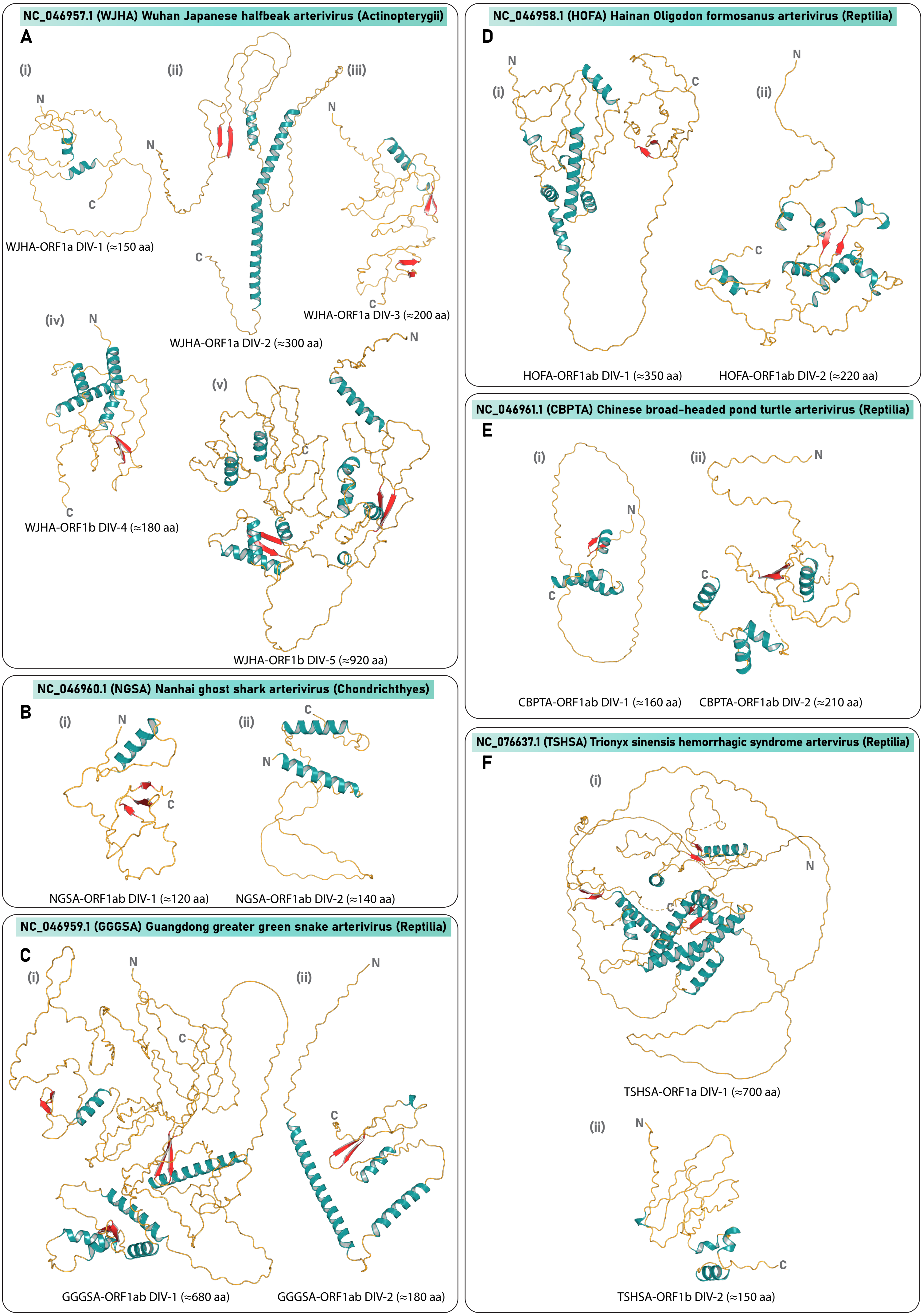

### Supplementary Figure S2

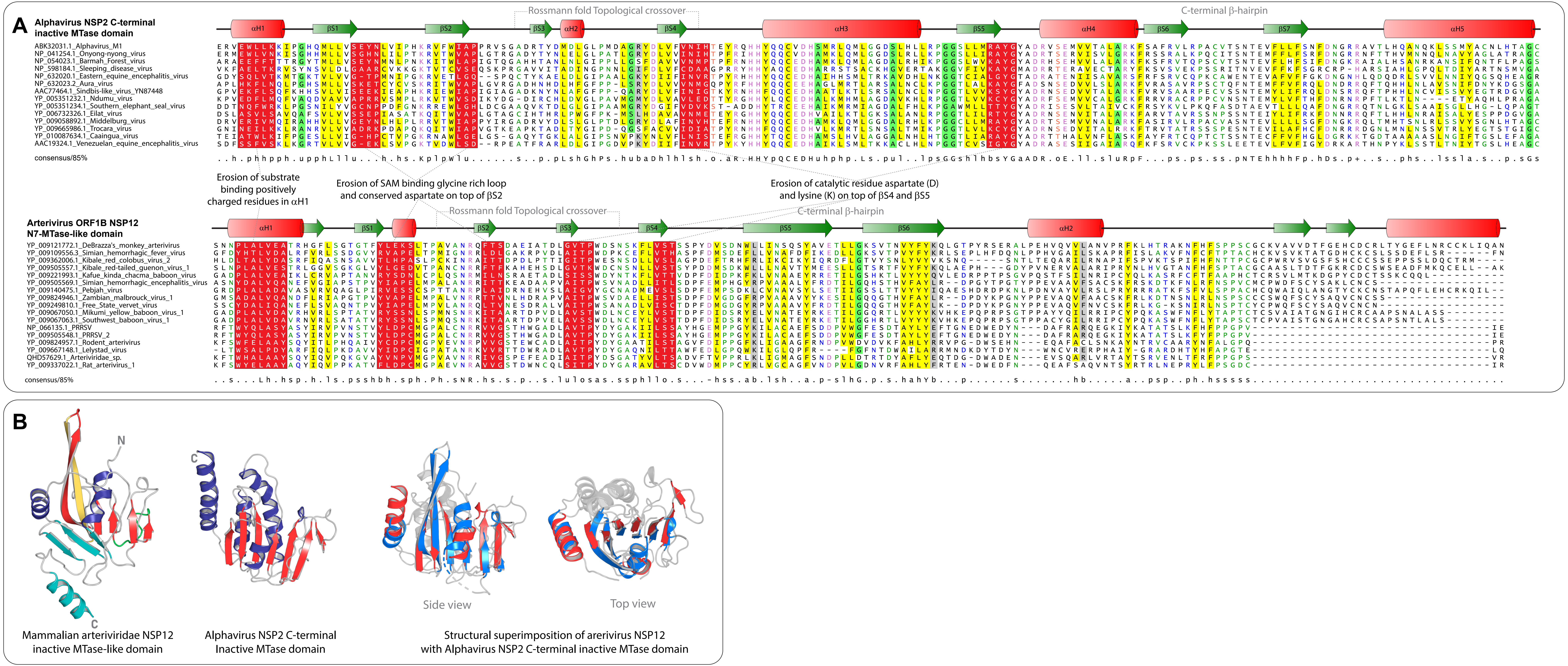

**Supplementary Figure S3**

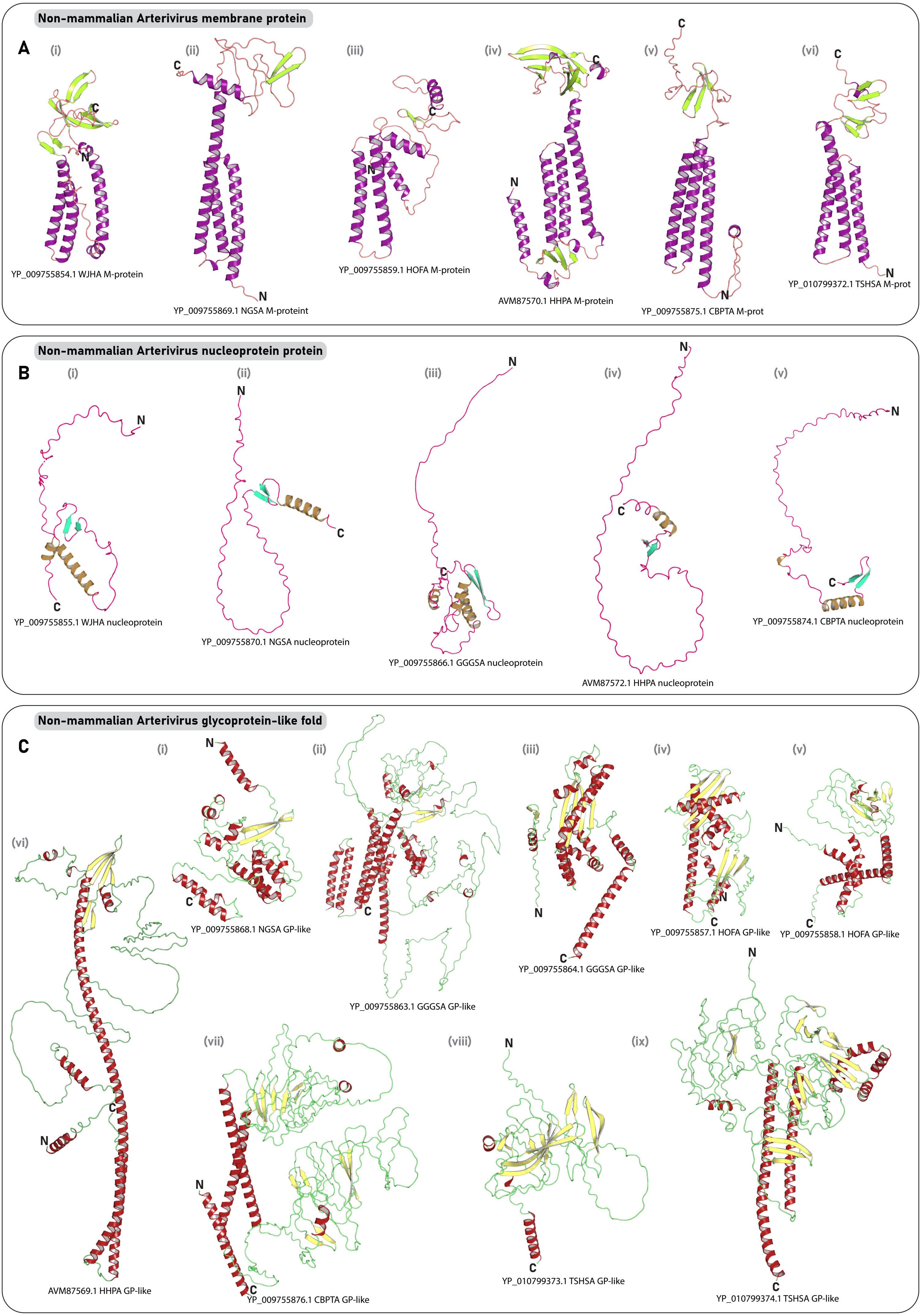

### Supplementary Figure S4

**E protein Tree Raw file Newick format**

(E_Prot_QHD57630.1_Bank_Voles_Rodent:1.1168901746,((((((((E_Prot_YP_009824950.1_Malbrouck_Primate:0.063
8362698,(E_Prot_YP_009505573.1_Simian_Primate:0.2608693601,E_Prot_YP_009249814.1_Chlorocebus_Primate:0.
0716678022)49/49:0.0883100614)97.4/81:0.3414677824,E_Prot_YP_009344811.1_Colobus_Primate:0.7179999757)7
8.8/25:0.1169980874,((E_Prot_YP_009121778.1_Monkey_Primate:0.0000025053,E_Prot_YP_009130634.1_Possum_
Marsupial:3.1878497073)59.7/38:0.2703608325,E_Prot_YP_009037600.1_Simian_Primate:0.0986674013)91.5/34:0.1
373672648)60.9/7:0.0646164667,(E_Prot_YP_009362010.1_Colobus_Primate:0.5195574072,E_Prot_YP_009140482.
1_Macaques_Primate:0.2289210928)79.6/35:0.0735366881)75/14:0.1114037388,(E_Prot_YP_009221999.1_Chicma_
Baboon_Primate:0.0026488989,(E_Prot_YP_009067069.1_Baboon_Primate:0.0663916246,E_Prot_YP_009067055.1_
Baboon_Primate:0.0390452073)99.8/92:0.5546817485)100/76:0.8806829909)66/24:0.0628968120,E_Prot_YP_00950
5561.1_Guenon_Primate:0.3725330472)99.3/75:1.0436069627,(E_Prot_YP_009388592.1_Shrew_Rodent:0.63797462
11,E_Prot_NP_127508.1_Equine_Horse:1.5776638580)30.8/40:0.2393085162)92.4/52:0.6090453293,E_Prot_YP_009
118962.1_Pouched_rat_Rodent:0.8054905197)79.5/51:0.3629167082,(((E_Prot_YP_009505550.1_Pig_Porcine:0.0000
010000,E_Prot_YP_006589918.1_Pig_Porcine:0.0149720218)98.7/97:0.2390188766,E_Prot_YP_009824959.1_Roden
t:0.1528902258)0/66:0.0163509188,E_Prot_YP_009337024.1_Rat_Rodent:0.1749661065)88.4/80:0.3697178514)84.2/
60:0.3123107765,E_Prot_YP_006589917.1_Env_Lactate:0.1657893621);

**GP2 protein Tree Raw file Newick format**

(GP2_NP_042574.1_Env_Lactate_dehydrogenase_elevating:0.6115135707,((GP2_NP_047408.1_Pig_Porcine:0.0000
010155,GP2_YP_009505549.1_Pig_Porcine:0.0086575404)89.4/98:0.0787461369,((GP2_YP_009337025.1_Rat_Rod
ent:0.1107882673,GP2_YP_009667149.1_Pig_Porcine:0.1407830999)70.9/72:0.0511030909,GP2_YP_009824958.1_
Rodent:0.0811980033)98.8/98:0.1874197223)100/96:0.4058973878,((((GP2_NP_065656.1_Equine_Horse:1.31026940
98,GP2_YP_009388593.1_Shrew_Rodent:1.7409437886)4.9/19:0.1310417811,((((((GP2_NP_203547.1_Simian_Prima
te:0.2708119761,GP2_YP_009121777.1_Monkey_Primate:0.3217990867)97.6/87:0.3355585014,GP2_YP_009140483
.1_Macaques_Primate:0.6416159515)93.8/58:0.3144353559,(((GP2_YP_009249815.1_Chlorocebus_Primate:0.11300
56635,GP2_YP_009824951.1_Malbrouck_Primate:0.2108730634)72/68:0.0996373017,GP2_YP_009505574.1_Simian
_Primate:0.2067571782)100/100:0.7378023206,GP2_YP_009362011.1_Colobus_Primate:0.6715488429)21.1/28:0.18
19370799)10.9/19:0.1023902565,GP2_YP_009344812.1_Colobus_Primate:0.8053150797)68.3/35:0.1283155575,((((G
P2_YP_009067056.1_Baboon_Primate:0.2459831780,GP2_YP_009067070.1_Baboon_Primate:0.1496928884)94.9/9
4:0.3780108268,GP2_YP_009222000.1_Chicma_Baboon_Primate:0.5598893876)96.6/99:0.6425606832,GP2_YP_00
9130633.1_Possum_Marsupial:1.4069307266)14.9/45:0.0773837302,GP2_YP_009505562.1_Guenon_Primate:1.0552
352378)95/74:0.4896808996)92.5/65:0.3269623989,((((((GP2__YP_009824947.1_Malbrouck_Primate:0.1205699366,
GP2__YP_009249811.1_Chlorocebus_Primate:0.1474435164)75.5/66:0.0465209760,GP2__YP_009505570.1_Simian
_Primate:0.1574782851)100/100:0.6133482657,(GP2__YP_009121774.1_Monkey_Primate:0.3421792283,GP2__NP_
203544.1_Simian_Primate:0.4707328394)8.6/30:0.0374251323)89.2/79:0.1186509944,GP2__YP_009505558.1_Guen
on_Primate:0.6702867385)91.3/79:0.2243591915,((GP2__YP_009362007.1_Colobus_Primate:0.3880745339,(GP2__
YP_009221996.1_Chicma_Baboon_Primate:0.1400907299,(GP2__YP_009067065.1_Baboon_Primate:0.0596684704,
GP2__YP_009067052.1_Baboon_Primate:0.0375178537)93.9/100:0.0812047512)100/100:0.4172682185)90.7/86:0.19
66114419,GP2__YP_009140478.1_Macaques_Primate:0.9624516589)89.2/75:0.1772954142)86/75:0.1919796351,GP
2__YP_009344808.1_Colobus_Primate:0.4992771787)99.9/97:0.7020170957)72/43:0.1535290071)97/69:0.49820055
57,GP2_YP_009118963.1_Pouched_rat_Rodent:1.0822355463)41/40:0.0823886803,GP2_QHD57631.1_Bank_Voles_
Rodent:1.1258820029)67.9/37:0.1149232244);

**GP3 protein Tree Raw file Newick format**

(GP3_YP_009824952.1_Malbrouck_Primate:0.3813600850,(((((((((GP3_YP_009667150.1_Pig_Porcine:0.1517091173,GP3_YP_009824960.1_Rodent:0.1920022899)41.1/39:0.0414548877,GP3_YP_009337026.1_Rat_Rodent:0.0688105246)88.6/54:0.0991512179,(GP3_YP_009505551.1_Pig_Porcine:0.0000020058,GP3_NP_047409.1_Pig_Porcine:0.0247261978)98.1/97:0.2194112649)99/92:0.4543521786,GP3_NP_042575.1_Env_Lactate:0.8553583087)86.6/62:0.2456134530,(GP3_YP_009118964.1_Pouched_rat_Rodent:1.0503711093,GP3_QHD57632.1_Bank_Voles_Rodent:0.7509672083)72/31:0.1229862720)88.5/51:0.3671362613,GP3_YP_009388594.1_Shrew_Rodent:2.0144204831)64.2/47:0.1659845643,(((((GP3_dup_YP_009824948.1_Malbrouck_Primate:0.1624730601,(GP3_dup_YP_009505571.1_Simian_Primate:0.1549919629,GP3_dup_YP_009249812.1_Chlorocebus_Primate:0.2087539616)82.8/82:0.0434716807)99.9/99:0.4394844519,GP3_dup_NP_203545.1_Simian_Primate:0.6294252035)77.1/40:0.0923757051,(GP3_dup_YP_009505559.1_Guenon_Primate:0.5107154295,GP3_dup_YP_009121775.1_Monkey_Primate:0.3903415170)99/97:0.3592407651)56.5/46:0.1793694895,((GP3_dup_YP_009362008.1_Colobus_Primate:0.4050773053,((GP3_dup_YP_009221997.1_Chicma_Baboon_Primate:0.3501583181,GP3_dup_YP_009067067.1_Baboon_Primate:0.1333356145)21.1/53:0.0481274320,GP3_dup_YP_009067053.1_Baboon_Primate:0.0504083492)100/100:0.6265306542)100/97:0.4782059673,GP3_dup_YP_009344809.1_Colobus_Primate:0.6876811005)13.1/23:0.0948806406)74.6/36:0.2691938117,GP3_dup_YP_009140480.1_Macaques_Primate:0.8644131593)96.9/94:0.6549423608)95.8/62:0.5252847640,((((GP3_YP_009505563.1_Guenon_Primate:1.3338873304,GP3_NP_065657.1_Equine_Horse:1.5447913286)0/10:0.0000024338,(GP3_YP_009362012.1_Colobus_Primate:0.7505485418,GP3_YP_009130635.1_Possum_Marsupial:1.9578578366)75.2/26:0.2272940704)70/9:0.1208112736,((GP3_YP_009121779.1_Monkey_Primate:0.7394529370,GP3_NP_203548.1_Simian_Primate:0.6823475683)96.3/82:0.4859309899,GP3_YP_009140484.1_Macaques_Primate:1.0402240189)52.9/28:0.2503291298)76.9/8:0.2331795496,(GP3_YP_009344813.1_Colobus_Primate:1.5147733220,GP3_YP_009222001.1_Chicma_Baboon_Primate:1.2481638114)57.4/18:0.2319025773)0/10:0.0000020341)72.8/37:0.3513476856,(GP3_YP_009067071.1_Baboon_Primate:0.4052642523,GP3_YP_009067057.1_Baboon_Primate:0.1073516108)99.8/100:1.2861789618)98/100:0.9343440223,(GP3_YP_009505575.1_Simian_Primate:0.3329408156,GP3_YP_009249816.1_Chlorocebus_Primate:0.3832829825)90.3/90:0.2426885651);

**GP4 protein Tree Raw file Newick format**

(GP4_NP_042576.1_Env_Lactate:0.6715681276,((((((GP4_NP_047410.1_Pig_Porcine:0.0051850849,GP4_YP_009505552.1_Pig_Porcine:0.0052794129)100/98:0.2196215885,GP4_YP_009824961.1_Rodent:0.1178970572)72.9/29:0.0487634100,GP4_YP_009337027.1_Rat_Rodent:0.0836882446)97/41:0.1540954641,GP4_YP_009667151.1_Pig_Porcine:0.0441660939)98.9/93:0.5545045275,GP4_QHD57633.1_Bank_Voles_Rodent:1.1333123401)18/31:0.1406585803,((((((GP4_NP_065658.1_Equine_Horse:2.8118628577,(((GP4_YP_009067058.1_Baboon_Primate:0.1646241313,GP4_YP_009067072.1_Baboon_Primate:0.1615791924)92.5/88:0.3411840938,GP4_YP_009222002.1_Chicma_Baboon_Primate:0.6928668442)78.2/90:0.4233104311,GP4_YP_009130636.1_Possum_Marsupial:3.1417946261)73/46:0.3707722054)73.2/20:0.3340618717,GP4_YP_009505564.1_Guenon_Primate:1.0032093386)80.1/13:0.3199290967,((GP4_NP_203549.1_Simian_Primate:0.5906027343,GP4_YP_009121780.1_Monkey_Primate:0.5228074190)98.8/94:0.6162699089,GP4_YP_009140485.1_Macaques_Primate:0.8290259999)86.9/42:0.2107071364)29/7:0.0851173931,(((GP4_YP_009249817.1_Chlorocebus_Primate:0.3901968758,GP4_YP_009505576.1_Simian_Primate:0.2525422823)100/98:1.1865003135,GP4_YP_009344814.1_Colobus_Primate:0.9377953963)83.8/37:0.2635683981,GP4_YP_009362013.1_Colobus_Primate:1.0309494234)19.2/14:0.1265443078)93.4/47:0.4565822500,(((((GP4_dup_NP_203546.1_Simian_Primate:0.5420657090,(GP4_dup_YP_009121776.1_Monkey_Primate:0.1641961648,GP4_dup_YP_009505560.1_Guenon_Primate:0.7278710367)88.2/76:0.1599913538)90/81:0.1853733224,(GP4_dup_YP_009249813.1_Chlorocebus_Primate:0.1639473624,(GP4_dup_YP_009505572.1_Simian_Primate:0.1784423670,GP4_dup_YP_009824949.1_Malbrouck_Primate:0.1644790122)42/39:0.0700186876)99.4/84:0.4599404030)95.4/87:0.3816032256,(((GP4_dup_YP_009067054.1_Baboon_Primate:0.1460796370,GP4_dup_YP_009067068.1_Baboon_Primate:0.1142137531)96.8/92:0.2309773099,GP4_dup_YP_009221998.1_Chicma_Baboon_Primate:0.1287066145)99.5/98:0.5341804554,GP4_dup_YP_009362009.1_Colobus_Primate:0.3882846402)97/96:0.4112017574)91.9/74:0.2688773500,GP4_dup_YP_009140481.1_Macaques_Primate:0.9696336333)0/34:0.0702718140,GP4_dup_YP_009344810.1_Colobus_Primate:0.6823996911)94.5/76:0.4502045753)97.4/63:0.6941135857,GP4_YP_009118965.1_Pouched_rat_Rodent:1.0853084663)80.2/47:0.3318350251)83.1/51:0.2800411546,GP4_YP_009388595.1_Shrew_Rodent:2.1689438591);

**GP5 protein Tree Raw file Newick format**

(GP5_YP_009824962.1_Rodent:0.3469831057,(((((((((((GP5_YP_009824954.1_Malbrouck_Primate:0.1500968002,G
P5_YP_009249819.1_Chlorocebus_Primate:0.1421421050)91.2/72:0.0929121112,GP5_YP_009505578.1_Simian_Pri
mate:0.1941820029)100/100:0.5255670959,GP5_YP_009362015.1_Colobus_Primate:0.5517001771)85.9/21:0.09448
39427,((GP5_YP_009505565.1_Guenon_Primate:0.6605938140,GP5_YP_009121781.1_Monkey_Primate:0.29785293
30)97.2/63:0.1882649017,((GP5_YP_009344816.1_Colobus_Primate:0.4324856663,GP5_NP_203550.2_Simian_Prim
ate:0.5080076637)87.1/50:0.1198772254,(GP5_YP_009140487.1_Macaques_Primate:0.4642038375,GP5_YP_00913
0637.1_Possum_Marsupial:3.1172733707)0/17:0.0000027624)64.6/6:0.0263520445)79.6/11:0.0424610427)46.2/28:0.
1241222735,(GP5_YP_009222003.1_Chicma_Baboon_Primate:0.4975801071,(GP5_YP_009067073.1_Baboon_Prim
ate:0.1687670135,GP5_YP_009067059.1_Baboon_Primate:0.2728490750)87.4/79:0.1055832727)80.2/79:0.11358271
92)95.8/92:1.0505394041,GP5_NP_065659.1_Equine_Horse:4.4919151360)72.2/52:0.4815035538,GP5_YP_0091189
66.1_Pouched_rat_Rodent:1.6972169520)43.9/32:0.1858522654,GP5_YP_009388596.1_Shrew_Rodent:2.746129355
7)91.8/63:0.5220736194,GP5_NP_042577.1_Env_Lactate_dehydrogenase_elevating:0.6927770022)69.6/35:0.365309
0153,GP5_YP_009667152.1_Pig_Porcine:0.5958627707)85/35:0.1192859612,GP5_YP_009337029.1_Rat_Rodent:0.2
797653108)88.4/42:0.1461849734,(GP5_YP_009505553.1_Pig_Porcine:0.0000010001,GP5_NP_047411.1_Pig_Porci
ne:0.0137052312)100/99:0.5255630701)7.7/36:0.0321322705,GP5_QHD57634.1_Bank_Voles_Rodent:0.2087132805)
;

**GP5a protein Tree Raw file Newick format**

(GP5a_YP_009824953.1_Malbrouck_Primate:0.0000029837,(GP5a_YP_009505577.1_Simian_Primate:0.2003239069,
GP5a_YP_009249818.1_Chlorocebus_Primate:0.5842329891)93.7/87:0.2956142289,((((((GP5a_YP_009388597.1_Sh
rew_Rodent:0.0000029086,GP5a_YP_006390077.1_Equine_Horse:3.4541536576)82.7/43:0.9682190670,((((GP5a_Y
P_009337028.1_Rat_Rodent:0.2320188741,GP5a_QHD57635.1_Bank_Voles_Rodent:0.2811182833)26.3/23:0.05550
90403,GP5a_YP_006488613.1_Pig_Porcine:0.6198720470)45.5/48:0.3701378927,GP5a_YP_009130638.1_Possum_
Marsupial:3.1434604456)62.3/36:0.7121552407,GP5a_YP_006488612.1_Env_Lactate_dehydrogenase_elevating:0.39
97876577)89.1/37:0.8835361713)92.3/42:1.4508686345,GP5a_YP_009037601.1_Simian_Primate:0.4321303155)82.9
/13:0.6342296382,((((GP5a_YP_009222004.1_Chicma_Baboon_Primate:1.3596881695,GP5a_YP_009067074.1_Bab
oon_Primate:0.2421256399)60/5:0.1429876957,GP5a_YP_009067060.1_Baboon_Primate:0.1327634741)96.3/18:0.78
99868208,GP5a_YP_009121782.1_Monkey_Primate:0.5382655908)54.2/23:0.2906199682,GP5a_YP_009118967.1_P
ouched_rat_Rodent:2.1716553289)70.4/19:0.2993030154)74.4/18:0.2136480508,(GP5a_YP_009344815.1_Colobus_P
rimate:0.3483684139,GP5a_YP_009140486.1_Macaques_Primate:1.1255371748)79.6/29:0.2739554888)97.5/73:0.91
05194494,GP5a_YP_009362014.1_Colobus_Primate:0.0000021608)100/90:1.4954222993);

**M protein Tree Raw file Newick format**

(M_Prot_YP_009824963.1_Rodent:0.0470667933,((((((M_Prot_YP_009824955.1_Malbrouck_Primate:0.0352592312,M_Prot_YP_009249820.1_Chlorocebus_Primate:0.0615960058)78.8/72:0.0326139312,M_Prot_YP_009505579.1_Simian_Primate:0.0398617705)98.4/95:0.2050673240,((((((M_Prot_YP_009505566.1_Guenon_Primate:0.1941836528,M_Prot_YP_009121783.1_Monkey_Primate:0.0875891013)72/58:0.0717310894,M_Prot_NP_203551.1_Simian_Primate:0.1541586272)88.2/79:0.0698839469,(M_Prot_YP_009222005.1_Chicma_Baboon_Primate:0.1872685229,(M_Prot_YP_009067075.1_Baboon_Primate:0.0088399587,M_Prot_YP_009067061.1_Baboon_Primate:0.0437471804)98.4/97:0.1533722851)99.3/77:0.1775951018)81.7/21:0.0724450347,M_Prot_YP_009140488.1_Macaques_Primate:0.1759574798)76/9:0.0278063844,M_Prot_YP_009362016.1_Colobus_Primate:0.2147960998)84.6/13:0.0665418375,M_Prot_YP_009344817.1_Colobus_Primate:0.2142589803)52.7/16:0.0438335625)99.3/70:0.5941323321,((M_Prot_YP_009388598.1_Shrew_Rodent:1.0468465609,(((M_Prot_YP_009130639.1_Possum_Marsupial:1.5951111438,((M_Prot_non-mamma_YP_009755859.1_Reptilia-snake:1.5279709990,M_Prot_non-mamma_YP_009755875.1_Reptilia-Turtle:1.2522094871)71.1/65:0.5710597717,M_Prot_non-mamma_YP_010799372.1_Reptilia-Turtle:2.3696967385)78.6/65:0.3719155489)69.7/50:0.4602014595,(M_Prot_non-mamma_YP_009755869.1_Chondrichthyes:1.6481443013,M_Prot_non-mamma_AVM87570.1_Reptilia-snake:1.3541574595)95/88:0.9353863722)79.5/66:0.3838671466,M_Prot_non-mamma_YP_009755854.1_Actinopterygii:2.7510960643)73.3/43:0.3612645787)51.4/32:0.2693270800,(M_Prot_YP_009118968.1_Pouched_rat_Rodent:0.6933718517,M_Prot_NP_065660.1_Equine_Horse:2.1750727998)82.9/35:0.4159492137)80.3/52:0.1989496190)96.7/75:0.4630428783,M_Prot_NP_042578.1_Env_Lactate:0.3470879541)97.7/85:0.3723201213,M_Prot_YP_009667153.1_Pig_Porcine:0.1067019032)94.6/59:0.1046297922,(((M_Prot_YP_009505554.1_Pig_Porcine:0.0117574517,M_Prot_NP_047412.1_Pig_Porcine:0.0000021101)99.7/99:0.0771333261,M_Prot_YP_009337030.1_Rat_Rodent:0.0559641878)84.5/41:0.0244014583,M_Prot_QHD57636.1_Bank_Voles_Rodent:0.1138275212)29.1/22:0.0067921371);

**N protein Tree Raw file Newick format**

(N_Prot_QHD57637.1_Bank_Voles_Rodent:0.0685793465,((((((((((((N_Prot_YP_009824956.1_Malbrouck_Primate:0.2332787109,(N_Prot_YP_009505580.1_Simian_Primate:0.4727385383,N_Prot_YP_009249821.1_Chlorocebus_Primate:0.0852755037)52.3/48:0.0697354001)99.7/87:0.5614720773,N_Prot_YP_009362017.1_Colobus_Primate:0.4123529930)91.6/68:0.2440311669,N_Prot_YP_009344818.1_Colobus_Primate:0.6929041921)87.7/47:0.2191399047,((((N_Prot_YP_009222006.1_Chicma_Baboon_Primate:0.3494101714,N_Prot_YP_009067076.1_Baboon_Primate:0.0279327343)50.5/41:0.0658613207,N_Prot_YP_009067062.1_Baboon_Primate:0.0595214742)81.4/75:0.3093331484,N_Prot_YP_009130640.1_Possum_Marsupial:2.2953662476)86.6/47:0.3288192218,N_Prot_YP_009140489.1_Macaques_Primate:0.3547284247)89.9/35:0.2148438654)77.7/11:0.1311458059,(N_Prot_YP_009505567.1_Guenon_Primate:0.6025220907,N_Prot_YP_009121784.1_Monkey_Primate:0.2326331815)80.8/51:0.1560315277)66.8/38:0.4956937732,((N_Prot_non-mammalian_YP_009755855.1_Actinopterygii:1.3292223273,N_Prot_non-mammalian_YP_009755870.1_Chondrichthyes:0.6333756073)77.8/68:0.4481022550,N_Prot_non-mammalian_AVM87572.1_Reptilia-snake:0.8491863617)97.6/98:1.1355916408)0/39:0.0169296591,N_Prot_NP_065661.1_Equine_Horse:1.4413288222)86.1/40:0.2654526790,(N_Prot_YP_009388599.1_Shrew_Rodent:0.6579618582,(N_Prot_non-mammalian_YP_009755866.1_Reptilia-snake:1.3086899197,N_Prot_non-mammalian_YP_009755874.1_Reptilia-Turtle:1.0989107109)87.8/60:0.5566849603)47/26:0.3428818080)35.9/31:0.0989760374,N_Prot_YP_009118969.1_Pouched_rat_Rodent:0.9433323913)98.5/94:0.5440660071,N_Prot_NP_042579.1_Env_Lactate:0.4673039543)93.5/81:0.2861724979,N_Prot_YP_009667154.1_Pig_Porcine:0.2321133582)79.7/61:0.0989056249,(N_Prot_YP_009505555.1_Pig_Porcine:0.0000010002,N_Prot_NP_047413.1_Pig_Porcine:0.0000010002)98.6/100:0.1650295191)92.8/83:0.1015759778,N_Prot_YP_009337031.1_Rat_Rodent:0.1648344552)31.2/46:0.0363963419,N_Prot_YP_009824964.1_Rodent:0.1467610299);
